# Proximity labeling of the *Listeria monocytogenes* surface reveals pathogen control of a host deubiquitinase

**DOI:** 10.64898/2026.02.17.706400

**Authors:** Patrick J. Woida, Maisie W. Smith, Rebecca L. Lamason

**Affiliations:** Department of Biology, Massachusetts Institute of Technology, Cambridge, MA, USA

## Abstract

Intracellular pathogens must navigate the crowded cellular environment to establish infection. *Listeria monocytogenes* achieves this by recruiting host factors to its surface to hijack the host actin cytoskeleton for motility, form membrane protrusions, and spread from cell to cell. Although these types of *Listeria*-host interactions are critical for infection, systematic characterization of this interface has been limited. Here, we implement surface display of the promiscuous biotin ligase split-TurboID to profile host proteins recruited to the surface of *L. monocytogenes* during intracellular infection. This approach identified the host deubiquitinase CYLD as a protein selectively enriched at the pathogen surface. While CYLD promotes infection by suppressing autophagy and innate immunity in macrophages, how *L. monocytogenes* recruits and appropriates CYLD function in other cell types has remained unclear. We demonstrate that the E3 ligase RNF213 decorates bacterial poles with M1-linked linear ubiquitin, thereby redirecting CYLD to the bacterial surface. We further show ubiquitin is not sufficient to recruit CYLD but requires the *L. monocytogenes* secreted effector internalin C (InlC). Despite its presence at the bacterial surface, CYLD does not deubiquitinate bacteria or regulate autophagic bacterial clearance in infected epithelial cells. Instead, CYLD and InlC protect *L. monocytogenes* from IFN-γ- and RNF213-dependent restriction of cell-to-cell spread. Overall, our work profiling the bacterial surface-host interactome has identified a new mechanism by which InlC spatially reprograms CYLD activity, uncoupling its canonical immune functions to promote cell-to-cell spread in epithelial cells. These findings highlight how *L. monocytogenes* exploits, a host deubiquitinase, to perform cell-type-specific functions during infection.

## INTRODUCTION

*Listeria monocytogenes* is a foodborne Gram-positive bacterial that can cause gastroenteritis in healthy individuals. In severe cases, *L. monocytogenes* can cross the intestinal barrier and enter the bloodstream, causing bacterial sepsis. From there, it may invade the liver and spleen or cross the blood–brain and placental barriers, leading to meningitis and abortion, respectively (1). A key attribute of *L. monocytogenes* that allows it to cause severe disease is its ability to invade, replicate, and disseminate within diverse cell types at various sites of infection, which calls for a comprehensive understanding of both conserved and cell-type-specific activities.

*L. monocytogenes* promotes its uptake into both phagocytic and non-phagocytic cells and escapes into the host cell cytosol. Once inside, it hijacks the host cytoskeleton to propel itself through the cytosol and eventually disseminates throughout host tissues via cell-to-cell spread (1, 2). Each step of the intracellular lifecycle requires direct interactions with host factors, navigation through the crowded cytosol, and remodeling of host cell membranes to invade or spread from cell to cell. *L. monocytogenes* directs some of these activities by recruiting and controlling host proteins at the bacterial surface. For example, *L. monocytogenes* expresses the membrane protein ActA to recruit the host Arp2/3 complex, VASP, and other actin regulators to the bacterial surface, thereby inducing polymerization of the actin tails that enable actin-based motility (2–4). *L. monocytogenes* can also recruit the host kinase CK2 to the bacterial surface to phosphorylate ActA to increase its binding affinity for the Arp2/3 complex and enhance actin-based motility (5).

In addition to enabling motility, surface-associated host factors can also protect *L. monocytogenes* inside the host cell. For example, actin and cytoskeletal regulators shield *L. monocytogenes* from autophagic recognition (6, 7). In their absence, host E3 ligases – including PARKIN and RNF166 – ubiquitinate the surface of *L. monocytogenes*, thereby recruiting p62 and other autophagy-related proteins to the bacterium and promoting its clearance through autophagy (8, 9). *L. monocytogenes* also uses the surface protein internalin K (InlK) to recruit the mammalian major vault protein to its surface to further camouflage the pathogen from autophagy (10). However, InlK expression has only been observed in mouse models of infection and can potentially vary across strains (7, 10), indicating that *L. monocytogenes* surface-associated host factors may vary across strains and intracellular environments.

These prior studies have established the critical paradigm that *L. monocytogenes* recruits various host factors to its surface to promote infection, but a comprehensive identification and understanding of these factors have been lacking because of our inability to broadly profile this space. Previously, sulfo-NHS-LC-biotin was used to non-specifically biotinylate the surface of bacterial pathogens (11–13). However, this labeling method requires lysis of host cells prior to labeling intracellular pathogens, which enriches for bacterial surface proteins rather than host-interacting proteins (12, 13). To overcome this hurdle, recent work successfully precoated *Shigella flexneri* in a purified peroxidase (APEX2) prior to intracellular invasion to enable biotinylation of host factors at the pathogen surface (14).Though the stability of surface localized recombinant APEX2 during long term infection has not been examined. To enable stable surface association, fusions between APEX2 and autotransporters have been proposed (15), but these have not been tested in intracellular infections and are restricted to Gram-negative bacteria. Thus, new approaches are needed to profile how Gram-positive bacteria like *L. monocytogenes* dynamically recruit and change surface-associated host factors to promote infection.

Here, we adapted split-TurboID to profile host proteins recruited to the *L. monocytogenes* surface during intracellular infection. We found that *L. monocytogenes* employs the secreted virulence factor InlC to recruit the mammalian deubiquitinase CYLD to sites of RNF213-dependent M1-linked ubiquitination at the bacterial poles. While CYLD has previously been shown to protect *L. monocytogenes* from autophagic clearance in macrophages, we show that InlC and CYLD promote cell-to-cell spread in epithelial cells. Altogether, our work profiling the host-bacterial surface interaction uncovers a new mechanism by which *L. monocytogenes* co-opts CYLD to promote infection.

## RESULTS

### Design of a surface-specific labeling tool for profiling *L. monocytogenes* interactions during infection

We sought to develop a proximity-labeling system that can be expressed on the bacterial surface during intracellular infection. We selected split-TurboID, which uses inactive fragments of the biotin ligase TurboID that only regain labeling activity when brought together in space (16). The advantage of this system is that it avoids potential deleterious effects of expressing a large heterologous protein cargo on the bacterial surface while maintaining flexibility for contact-dependent studies. To test the compatibility of this system with *L. monocytogenes*, we expressed the N-terminal domain of TurboID (NTurboID) fused to an HA epitope on the bacterial surface and we expressed the C-terminal domain (CTurboID) fused to a V5 epitope in the host cell cytosol. Once *L. monocytogenes* invades the host cell, the two halves of split-TurboID will reform to biotinylate proteins at the bacterial surface (**Fig. 1A**). To localize NTurboID-HA to the *L. monocytogenes* surface, we generated a panel of surface display constructs adapted from known *L. monocytogenes* surface proteins. Each were placed under the control of the *actA* promoter to restrict expression to the host cytosol and were integrated into the genome of *L. monocytogenes* (17) (**Table S1**). We found that fusions of NTurboID-HA to the internalin A (InlA) or internalin F (InlF) LPXTG-containing C-terminal surface-anchoring domains failed to support mammalian cell infection, suggesting these fusions were toxic when expressed. Similarly, fusing NTurboID-HA to the *L. monocytogenes* ActA transmembrane region did not support intracellular infection (**Table S1**).

**Figure 1.**
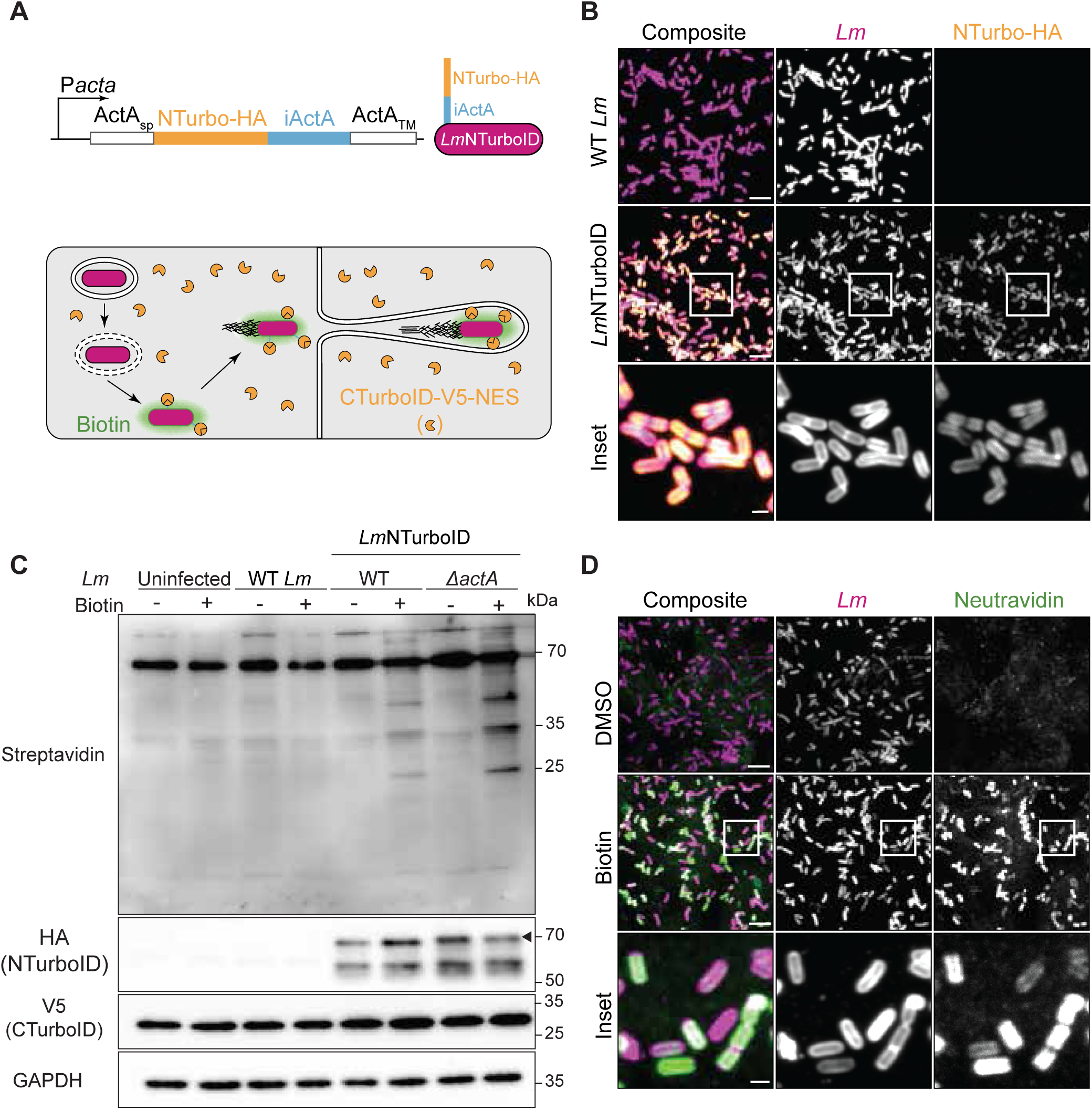
Split-TurboID allows for biotinylation at the bacterial-host surface during *L. monocytogenes* intracellular infection. **(A)** Schematic of surface display construct and application of split-TurboID. A fusion of the *L. monocytogenes* ActA signal peptide (ActA_sp_) and ActA transmembrane region, flanking NTurboID-HA and residues 336-843 of *L. ivanovii* ActA (iActA), was expressed under the control of the *actA* promoter. This construct was integrated into the *L. monocytogenes* genome to generate *Lm*NTurboID. Infection of cells stably expressing the cytosol-localized CTurboID (CTurboID-V5-NES) with *Lm*NTurboID reconstitutes TurboID activity, enabling biotinylation of proteins at the bacterial surface. (**B**) Immunofluorescence detection of NTurboID-HA localizing around bacteria in infected A549 cells at 5 hpi. NTurboID-HA, anti-HA (orange); *Lm, L. monocytogenes* antisera (magenta). (**C**) Western blot analysis of split-TurboID activity in A549-CTurboID cells infected with indicated strains and treated with DMSO or 50 µM biotin. Arrowhead indicates the 69 kDa full-length NTurboID-HA surface display construct. Data are representative of three independent experiments. (**D**) Immunofluorescence detection of split-TurboID activity in LmNTurboID-infected A549-CTurboID cells treated with DMSO or 50 µM biotin. Neutravidin-mediated detection of biotinylated proteins (green); *Lm, L. monocytogenes* antisera (magenta). **(B**, **D**) Images are representative of three independent experiments. Scale bar, 5 µm; inset scale bar, 1 µm.

We next considered whether expressing NTurboID too close to the bacterial cell wall was deleterious. Membrane-bound surface proteins of Gram-positive bacteria require long, highly disordered internal domains to provide the protein with significant free energy to extend the protein beyond the cell wall (18). *L. ivanovii* ActA (iActA) contains an internal disordered linker region that is longer than *L. monocytogenes* ActA (19), which we reasoned may be sufficient to extend heterologous cargo like NTurboID beyond the cell wall. Therefore, we modified the membrane-bound construct to include the iActA linker residues, 336-843, between NTurboID-HA and the *L. monocytogenes* ActA transmembrane domain. This final construct was integrated into the genome of *L. monocytogenes* as above and the strain designated as *Lm*NTurboID (**Fig. 1A**). To confirm NTurboID localization to the bacterial surface, we infected A549 cells with this modified strain and examined HA localization by immunofluorescence microscopy. *Lm*NTurboID readily infected cells and NTurboID-HA was observed on the perimeter of this modified strain, and not the parental strain of *L. monocytogenes,* indicating minimal-toxic expression and successful surface localization of NTurboID (**Fig. 1B**).

To localize CTurboID to the host cell cytosol, we generated A549 cells stably expressing CTurboID-V5 fused to a nuclear export signal (NES). A549 cells have previously been used to study *L. monocytogenes* intracellular growth and cell-to-cell spread, and they provide a highly tractable system to study host-pathogen interactions in non-phagocytic cells (20). We infected these A549-CTurboID cells with *Lm*NTurboID and added either DMSO or exogenous biotin to confirm that the cytosolic CTurboID and bacterial-surface localized NTurboID can be functionally reconstituted and biotinylate proteins at the bacterial-host interface. Western blots from infected cell lysates showed that split-TurboID-dependent biotinylation only occurred in cells infected with *Lm*NTurboID treated with biotin and not from cells treated with wild-type *L. monocytogenes* (**Fig. 1C**). We further examined biotinylation by microscopy to confirm biotinylation was restricted to the bacterial surface. Biotin signal, as detected by staining with fluorescently-conjugated neutravidin, was restricted to the bacterial surface in A549-CTurboID cells infected with *Lm*NTurboID and treated with endogenous biotin and not DMSO (**Fig. 1D**). These results indicate that bacterial surface display of NTurboID and host cytosolic expression of CTurboID allow for site-specific biotinylation at the bacterial surface during intracellular infection.

### Split-TurboID identifies host proteins recruited to the bacterial surface during intracellular infection

We next tested if split-TurboID can be used to selectively enrich for host proteins at the bacterial surface by comparing wild type and an actin-based motility deficient (Δ*actA*) strain of *L. monocytogenes*, as these have been previously shown to differentially associate with regulators of the cytoskeleton and autophagy, respectively (3, 4, 6, 7). A549-CTurboID cells were infected for 6 h with *L. monocytogenes* strains expressing NTurboID in either a wild-type or non-motile Δ*actA* background. Infected cells were treated with exogenous biotin during the last 5 hours, and streptavidin enrichment from collected lysates was performed, followed by mass spectrometry. We identified a total of 2382 proteins, with 49 proteins significantly enriched in wild-type *Lm*NTurboID-infected cells and 91 proteins significantly enriched in Δ*actA Lm*NTurboID-infected cells (**Fig. 2A**, **Table S2**). ActA was only identified in wild-type *Lm*NTurboID-infected cells and thus serves as a positive control between the two conditions (**Fig. 2A**). p62 was also specifically detected in Δ*actA Lm*NTurboID-infected cells, which agrees with prior data showing stable association between p62 and non-motile bacteria to enable autophagy-mediated clearance (6, 7). STRING network and gene ontology analysis revealed proteins enriched from Δ*actA Lm*NTurboID-infected cells are associated with innate immune regulation, specifically proteins involved in autophagy (e.g., NAP1, TBK1, SPATA2, and SQSTM1), while proteins enriched from wild-type *Lm*NTurboID-infected cells were associated with cytoskeleton regulation (e.g., KIAA1671, MISP, MYO6, LIMA1, LASP1, and FGD6) (**Fig. 2A** and **B**). While some Arp2/3 complex family members, such as ARPC3, ARPC4, and ARPC5, along with Ena/VASP, were identified; they were not significantly enriched between wild-type and Δ*actA Lm*NTurboID*-*infected cells (**Table S2**), which may reflect the transient nature of their proximity to the pathogen surface or a limitation of the hybrid ActA-Split-TurboID fusion. Taken together, though, these results show that split-TurboID can enrich for host proteins that are differentially associated with the two strains.

**Figure 2.**
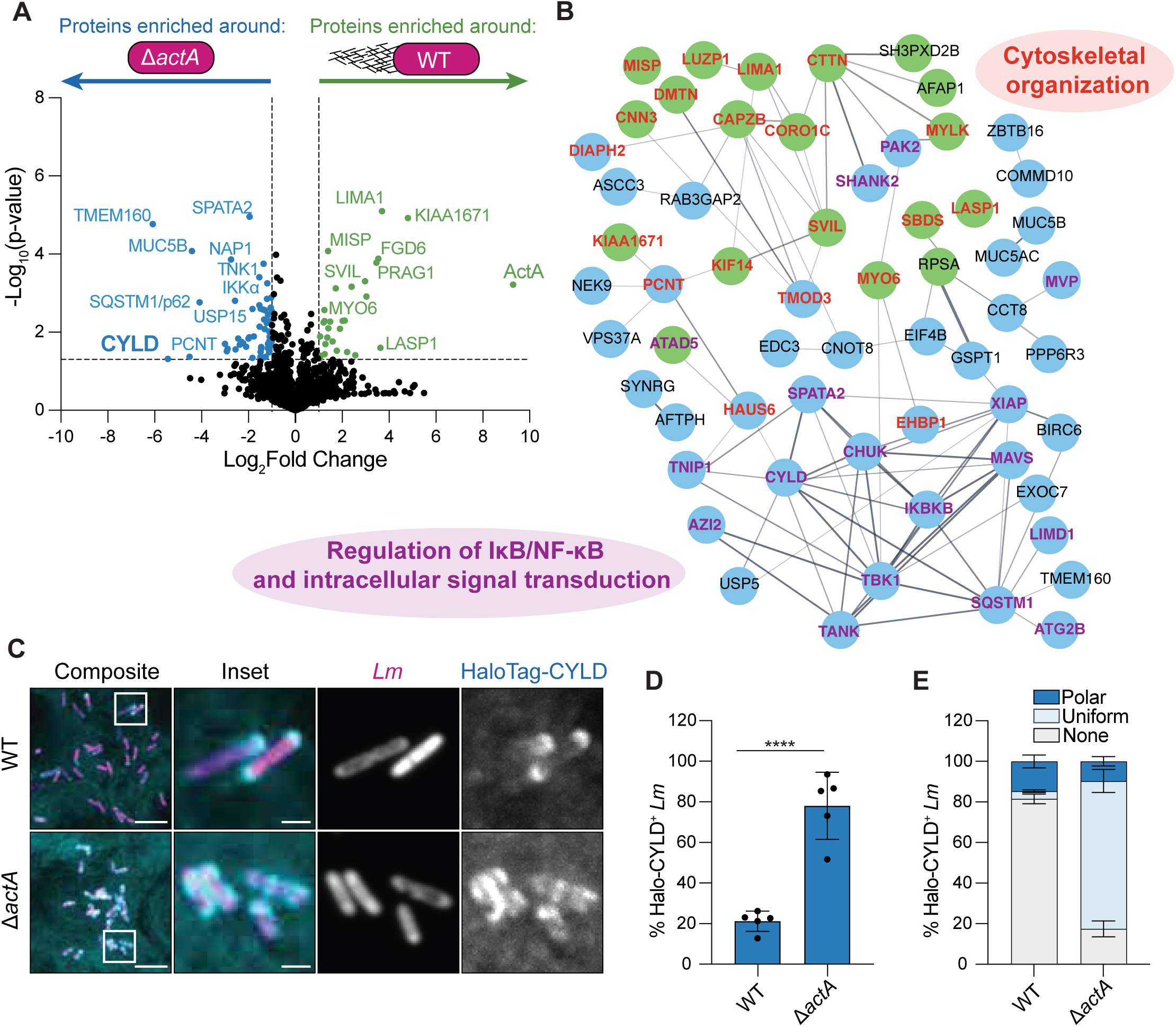
Proteomic profiling of the *L. monocytogenes* surface identifies the deubiquitinase CYLD localizes to *L. monocytogenes* during infection. **(A)** Volcano plot of proteins enriched from either wild-type or Δ*actA Lm*NTurboID infected A549-CTurboID cells. Dotted lines indicate cut-offs at -log_10_(p-value) > 1.3 (x-axis) and log_2_ fold change of 0.5 (y-axis). (**B**) STRING network analysis of significantly enriched proteins from wild-type (green) or Δ*actA* (blue) *Lm*NTurboID infected A549-CTurboID cells. Gene Ontology pathway analysis, along with manual curation, identified proteins in the network as regulators of cytoskeleton organization (GO: 0007010, red) along with IκB/NF-κB and intracellular signal transduction (GO:0043122, GO:1902531, purple). **(C)** Confocal microscopy of CYLD localization around indicated *L. monocytogenes* strains in infected A549-HaloTag-CYLD cells at 5 hpi. *Lm, L. monocytogenes* antisera (magenta); HaloTag-CYLD, HaloTag ligand Janelia Fluor 635 (cyan). Images are representative of three independent experiments. Scale bar, 5 µm; inset scale bar, 1 µm. (**D** and **E**) Percentage of total *Lm*Dasherco-localized with HaloTag-CYLD (**D**) or (**E**) co-localized at the poles, uniformly around the bacteria, or no observed co-localization at 5 hpi. Each data point represents the percentage of *Lm*Dasher per field of view (FOV). Data are representative of three independent experiments, with >250 bacteria counted per FOV in each experiment. Student’s t-test, ****P<0.0001.

Split-TurboID also identified host proteins that had been previously shown to regulate *L. monocytogenes* infection but not shown to localize to the pathogen surface, including the deubiquitinase CYLD and its regulator SPATA2, both of which were significantly enriched from Δ*actA* LmNTurboID*-*infected cells (**Fig. 2A, Table S2**). CYLD is a negative regulator of antibacterial immunity, and while prior work established that CYLD is necessary to promote *L. monocytogenes* infection in cultured macrophages and *in vivo* (21, 22). However, the mechanism by which *L. monocytogenes* co-opts CYLD and what role its localization plays during infection remainuntested. Therefore, we generated A549 cells stably expressing HaloTag-CYLD and infected them with either wild type or Δ*actA L. monocytogenes* strains expressing Dasher GFP (*Lm*Dasher) to validate CYLD localization. CYLD localized to both the wild type and Δ*actA Lm*Dasher but was significantly enriched around the Δ*actA* mutant, in agreement with our mass spectrometry data (**Fig. 2C** and **D**). Additionally, we observed divergent localization patterns between wild-type and Δ*actA Lm*Dasher. CYLD was primarily restricted at the poles of wild-type *Lm*Dasher, while CYLD localized uniformly around the surface of the Δ*actA* mutant (**Fig. 2E**). Thus, in addition to identifying host proteins known to localize to *L. monocytogenes*, we also validated a new host-pathogen interaction at the surface of *L. monocytogenes* during intracellular infection.

### CYLD is recruited to M1-linked ubiquitin at the bacterial poles

CYLD binds ubiquitinated substrates (23), suggesting it may be targeted to the pathogen via surface ubiquitination. Ubiquitin chains form through linking ubiquitin at various lysine residues or between the first N-terminal methionine residue and the C-terminal glycine residue to form linear M1-linked chains (24). Previous work showed that the Δ*actA* strain is heavily ubiquitinated (6), which may recruit CYLD, but it was unclear what was driving polar localization of CYLD in wild-type *L. monocytogenes*. We first examined polyubiquitin localization around both wild-type and Δ*actA Lm*Dasher by microscopy using the FK1 antibody, which preferentially binds to K29, K48, and K63 polyubiquitin linkages. As previously demonstrated, wild-type *Lm*Dasher is minimally ubiquitinated while the Δ*actA* mutant is robustly coated in polyubiquitin (**Fig. 3A** and **B**) (6, 7). While these data can account for CYLD enrichment around the Δ*actA* mutant, they fail to address how CYLD localizes to wild-type *Lm*Dasher. Therefore, we also examined M1-linked ubiquitin and observed that about ∼34% of wild-type *Lm*Dasher and almost all the Δ*actA* mutants were positive for M1-linked ubiquitin (**Fig. 3A** and **B**). Additionally, M1-linked ubiquitination displayed the same polar localization around wild-type *Lm*Dasher as CYLD (**Fig. 2C** and **E**, **Fig. 3A** and **C**). Thus, our data demonstrate that *L. monocytogenes* is targeted for M1-linked ubiquitination, which could drive recruitment of CYLD to the bacteria.

**Figure 3.**
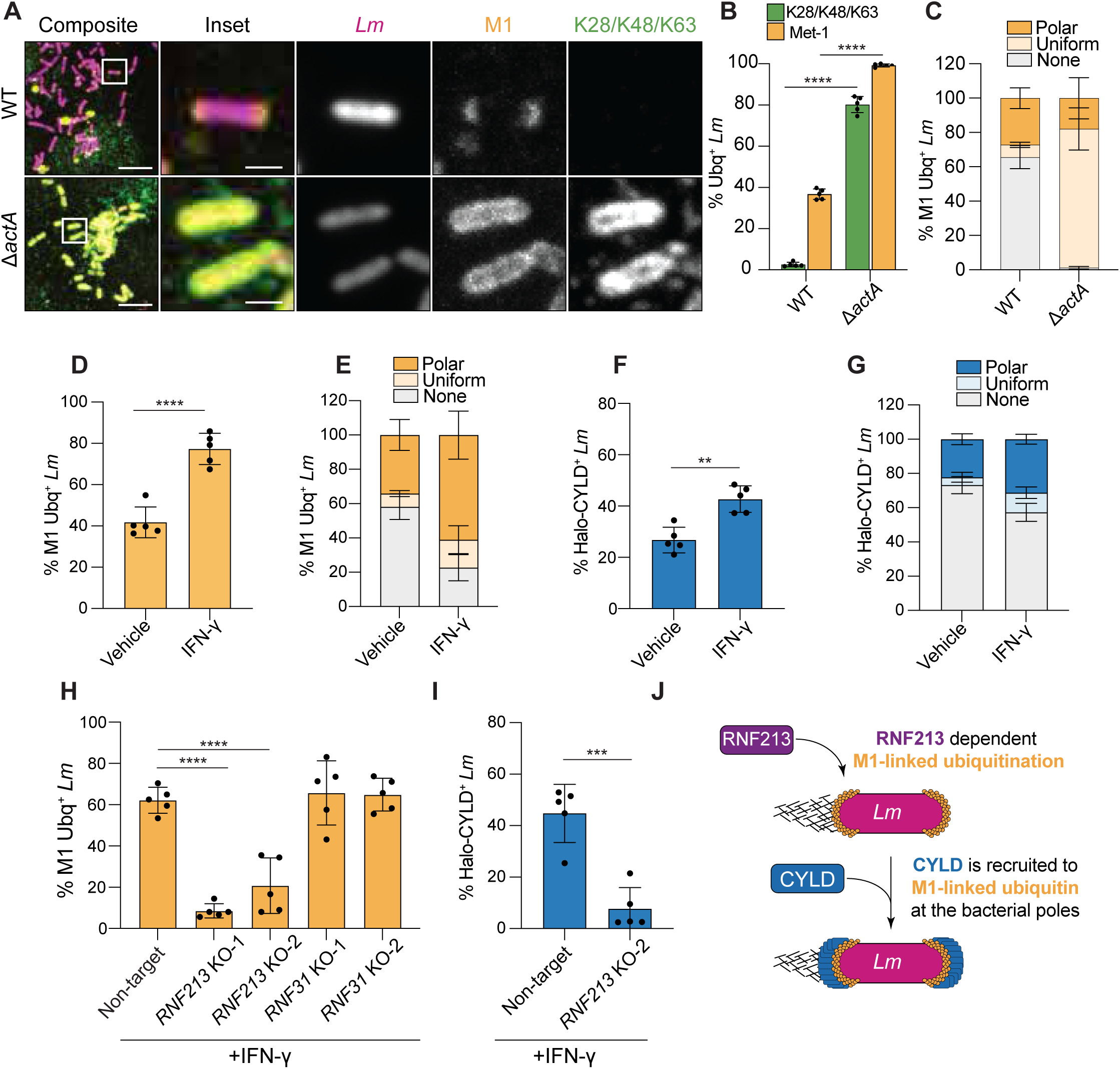
CYLD is recruited to RNF213-dependent M1-linked ubiquitination at the poles of *L. monocytogenes*. **(A)** Immunofluorescence detection of K29/K48/K63 polyubiquitin and M1-linked ubiquitin to indicated *Lm*Dasher strains at 5 hpi in A549 cells. *Lm, LmDasher* (magenta); anti-polyubiquitin (FK1 antibody, green); anti-M1-linked linear ubiquitin (orange). Images are representative of two independent experiments. Scale bar, 5 µm; inset scale bar, 1 µm. (**B** and **C**) Percentage of total wild-type and Δ*actA Lm*Dasher from (**A**) co-localized with K29/K48/K63 polyubiquitin and M1-linked ubiquitin (**B**), and with Met1-linked ubiquitin co-localized at the poles, uniformly around wild-type *Lm*Dasher, or no observed co-localization (**C**). Data are representative of two independent experiments. (**D** and **E**) Percentage of total wild-type *Lm*Dasher co-localized with M1-linked ubiquitin (**D**), and with M1-linked ubiquitin co-localized at the poles, uniformly around the bacteria, or no observed co-localization (**E**) at 5 hpi in infected A549 cells, with or without IFN-γ. Data are representative of two independent experiments. (**F** and **G**) Percentage of total *Lm*Dasher co-localized with HaloTag-CYLD (**F**), and with Halo-CYLD co-localized at the poles, uniformly around the bacteria, or no observed co-localization (**G**) at 5 hpi in A549 cells, with and without IFN-γ treatment. Data are representative three independent experiments. (**H**) Percentage of *Lm*Dasher co-localized with M1-linked ubiquitin in A549 cells with IFN-γ treatment at 5 hpi in *RNF213* or *RNF31* (HOIP) KO cells. Data representative of two independent experiments. Ordinary one-way ANOVA using Tukey’s multiple comparisons post-hoc test \*\*\**P*<0.00001. (**I**) Percentage of *Lm*Dasher co-localized with HaloTag-CYLD in *RNF213* KO cells. Data are representative of three independent experiments. (**J**) Model for RNF213-dependent, M1-linked ubiquitination of bacterial poles that leads to recruitment of CYLD to *L. monocytogenes.* All data are representative of the percentage of *Lm*Dasher per FOV displayed as mean ± SD; >150 bacteria were counted per FOV. For (**D**), (**F**), and (**I**) significance was determined using Student’s t-test\*\**P*<0.01. \*\*\**P*<0.00001.

CYLD protects against IFN-γ dependent autophagic clearance of *L. monocytogenes* in macrophages (22). IFN-γ also amplifies M1-linked ubiquitination around intracellular bacteria (25), suggesting IFN-γ may lead to enhanced localization of both M1-ubiquitin and CYLD on *L. monocytogenes*. To test this hypothesis, we pretreated A549 cells with IFN-γ for 24 h before infection with *Lm*Dasher and examined M1-linked linear ubiquitination by microscopy. IFN-γ significantly increased the percentage of M1-linked ubiquitin-positive *Lm*Dasher (**Fig. 3D**). This increase did not impact the distribution of M1-linked ubiquitin, as it remained predominantly localized to the bacterial poles (**Fig. 3E**). Pretreating A549-HaloTag-CYLD cells with IFN-γ before infection with *Lm*Dasher also increased the percentage of CYLD-positive *Lm*Dasher (**Fig. 3F**). Like M1-linked ubiquitin, IFN-γ did not impact HaloTag-CYLD distribution around *Lm*Dasher as it remained predominantly localized to the bacterial poles (**Fig. 3G**). These results show that IFN-γ enhances M1-linked ubiquitination and CYLD recruitment to the bacterial poles.

Intracellular bacterial pathogens are targeted by M1-linked ubiquitination through two mechanisms. The first occurs when bacteria are coated in K-linked polyubiquitin by E3 ligases (e.g., RNF213), which functions as a receptor for the linear ubiquitination chain assembly complex (LUBAC) that coats the pathogen in M1-linked ubiquitin (26–28). The second is through an RNF213-dependent, LUBAC-independent mechanism that is yet to be fully characterized (25). To determine the mechanism required for M1-linked ubiquitination of *L. monocytogenes,* we generated knockout (KO) cell lines using Cas12a with guides targeting either RNF213 or HOIP (RNF31), the latter of which is an essential subunit of the LUBAC complex (29) (**Fig. S1A** and **B**). We pretreated these cell lines, along with a cell line expressing a non-target guide, with IFN-γ for 24 h before infection with *Lm*Dasher. While M1-linked ubiquitination was observed in non-target infected cells, it was significantly reduced in both *RNF213* KO cell lines (**Fig. 3H**). In contrast, the percentage of M1-linked ubiquitin-positive *Lm*Dasher in the *RNF31* KO cells was comparable to non-target cells (**Fig. 3H**). These results show that *L. monocytogenes* is targeted by M1-linked ubiquitination in an RNF213-dependent and LUBAC-independent mechanism.

Because enhanced M1-linked ubiquitination correlated with an increase in CYLD localization to *L. monocytogenes* poles, we predicted that the loss of M1-linked ubiquitination would reduce CYLD recruitment. Therefore, we generated non-target control and *RNF213* KO cell lines stably expressing HaloTag-CYLD to test if CYLD localization to *L. monocytogenes* was dependent on M1-linked ubiquitination. We found that loss of *RNF213* expression dramatically attenuated HaloTag-CYLD recruitment to *Lm*Dasher (**Fig. 3I**), indicating that M1-linked ubiquitination is required for CYLD localization to the poles of intracellular *L. monocytogenes.* Taken together, our data suggests RNF213 ubiquitinates the poles of the *L. monocytogenes*, which further recruits CYLD to the bacteria (**Fig. 3J**).

### The secreted effector InlC enables selective recruitment of CYLD to ubiquitinated bacteria

CYLD has various ubiquitin-binding domains (23), prompting us to ask if its recruitment was specific to *L. monocytogenes* or found with any ubiquitinated bacterium targeted for autophagy. To test these possibilities, we examined CYLD localization around the obligate intracellular bacterial pathogen *Rickettsia parkeri*. *R. parkeri* normally avoids autophagy by methylating lysine residues on surface proteins, particularly the outer surface protein OmpB (30, 31). Transposon disruption of *ompB* leads to both K-linked and M1-linked ubiquitination of the *R. parkeri* surface and targets the pathogen for autophagic clearance (30, 31). We infected A549 HaloTag-CYLD cells with either wild-type *R. parkeri* or an *ompB* mutant (*ompB*^STOP^::Tn) and imaged CYLD localization. HaloTag-CYLD was not observed around either wild-type or the *ompB*^STOP^::Tn mutant (**Fig. 4A**). These data suggest that the M1-linked or K-linked ubiquitin that coats *R. parkeri* is not sufficient to accumulate CYLD at the bacterial surface.

**Figure 4.**
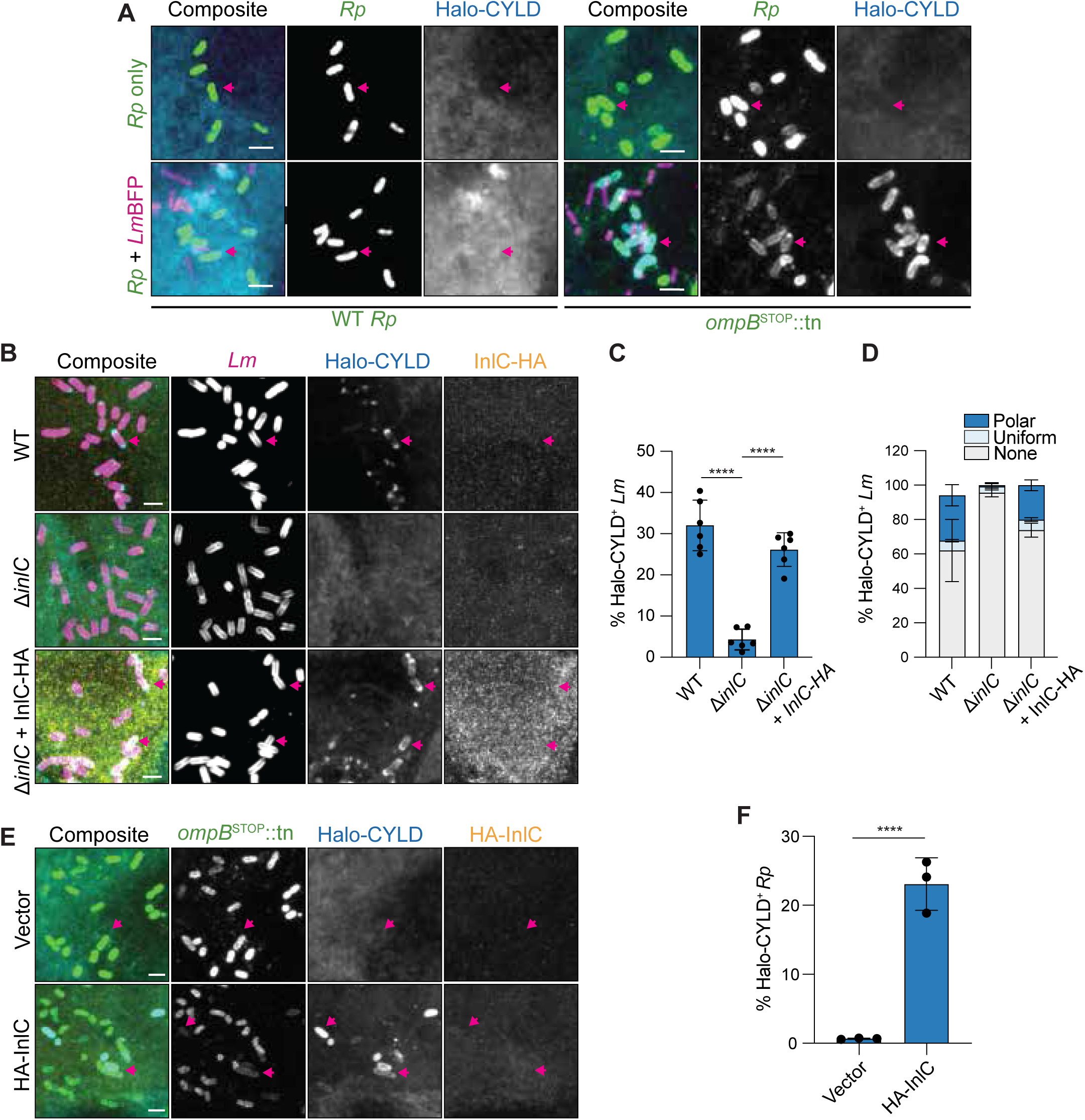
The *L. monocytogenes* secreted effector InlC re-localizes CYLD to ubiquitinated bacteria. **(A)** Immunofluorescence detection of HaloTag-CYLD localization in A549 cells upon infection with the indicated bacterial strains. *ompB*^Stop^::Tn *R. parkeri* mutant (Rp, green); BFP-expressing *L. monocytogenes* (*Lm*BFP, magenta); HaloTag-CYLD, HaloTag ligand (cyan). (**B**) Immunofluorescence detection of HaloTag-CYLD localization in A549 cells upon infection with the indicated *L. monocytogenes* strains at 5 hpi.*Lm*, *L. monocytogenes antisera* (magenta), anti-HA (HA-InlC, orange), HaloTag ligand Janelia Fluor 635 (cyan). (**C** and **D**) Percentage of indicated bacteria from (**B**) co-localized with HaloTag-CYLD or (**D**) with HaloTag-CYLD co-localized at the poles, uniformly around the bacteria, or no observed co-localization. Ordinary one-way ANOVA using Tukey’s multiple comparisons post-hoc test \*\*\**P*<0.00001. (**E**) Immunofluorescence detection of HaloTag-CYLD localization around the *ompB*^Stop^::Tn *R. parkeri* mutant in A549-HaloTag-CYLD cells stably transduced with an empty vector or one containing HA-InlC. Anti-rickettsia (*ompB*^Stop^::Tn *R. parkeri,* green), anti-HA (HA-InlC, orange), and HaloTag ligand Janelia Fluor 635 HaloTag Ligand (cyan). (**F**). Percentage of the *ompB*^Stop^::Tn *R. parkeri* mutant bacteria from (E) co-localized with HaloTag-CYLD at 5 hpi. Student’s t-test, ****P<0.00001. All immunofluorescence images and quantified data are representative of three independent experiments. White arrows indicate examples of bacteria with or without CYLD localization. Scale bar, 2 μm. All data are representative of the percentage of indicated bacteria per FOV displayed as mean ± SD; >250 bacteria were counted per FOV in each experiment.

We next investigated if a *L. monocytogenes-*specific factor was required for CYLD recruitment to ubiquitin-coated bacteria. We reasoned that *L. monocytogenes* may secrete a virulence factor (28–32) that modulates CYLD’s interaction with ubiquitinated substrates, or that CYLD may instead recognize a unique ubiquitinated substrate on the *L. monocytogenes* surface. To distinguish between these possibilities, we coinfected A549 HaloTag-CYLD cells with BFP-expressing *L. monocytogenes* (*Lm*BFP) and the wild-type and ubiquitin-rich *ompB*^STOP^::Tn mutant *R. parkeri* strains. If *L. monocytogenes* secretes a factor to induce CYLD localization to ubiquitin, *Lm*BFP should redirect CYLD to ubiquitinated *ompB*^STOP^::Tn mutant *R. parkeri.* However, if CYLD recognizes a *L. monocytogenes*-specific ubiquitinated substrate, CYLD should remain restricted to the poles of *Lm*BFP. We found that HaloTag-CYLD accumulated around the *ompB*^STOP^::Tn mutant only during coinfection with LmBFP, and not around wild-type *R. parkeri* (**Fig. 4A**). Therefore, we concluded that *L. monocytogenes* secretes a factor that induces CYLD localization to ubiquitinated substrates.

We sought to identify the *L. monocytogenes-*specific factor that controls CYLD localization. A prior high-throughput split-luciferase binding screen indicated a potential interaction between the multifunctional secreted bacterial protein internalin C (InlC) and CYLD (32). Therefore, we first assessed binding using AlphaFold3 and obtained a high confidence predicted interaction between InlC and CYLD (ipTM = 0.75) (**Fig. S2A** and **B**). We then tested if the proteins interacted during infection of epithelial cells. Using a HaloTag pulldown, we detected a potent interaction between HA-tagged InlC (expressed from Δ*inlC L. monocytogenes*) and HaloTag-CYLD, but not with HaloTag alone (**Fig. S2C**). We next tested if InlC was required for CYLD recruitment to bacteria. We found that deletion of *inlC* from *L. monocytogenes* significantly attenuated CYLD localization to the bacterial surface in A549-HaloTag-CYLD cells, and that polar CYLD localization was restored when HA-InlC was expressed in the Δ*inlC* background (**Fig. 4B - D**). We noted that HA-InlC did not co-localize with HaloTag-CYLD around *L. monocytogenes* (Δ*inlC* + HA-InlC complemented strain) (**Fig. 4B**) or *ompB*^STOP^::Tn *R. parkeri* (**Fig. 4E**). These results agree with previous data showing that endogenous InlC localizes diffusely throughout the host cytosol rather than around bacteria (33). To determine if InlC was sufficient to re-localize CYLD to ubiquitinated substrates, we generated HaloTag-CYLD A549 cells that also stably expressed HA-InlC. Using these cells, we observed CYLD localization around the *ompB*^STOP^::Tn mutant *R. parkeri* only in the presence of HA-InlC and not when infecting vector control cells (**Fig. 4E** and **F**). Thus, the secreted effector InlC is both necessary and sufficient to induce CYLD localization to ubiquitinated bacteria.

### CYLD does not protect *L. monocytogenes* from autophagy in epithelial cells

While CYLD protects *L. monocytogenes* from IFN-γ-induced autophagic bacterial clearance in macrophages (22), whether it plays a similar role in non-phagocytic cells remains unknown. Therefore, we asked whether CYLD protected *L. monocytogenes* from IFN-γ-induced autophagic clearance in epithelial cells. To test this hypothesis, we generated two *CYLD* knockout (KO) lines and examined whether CYLD protects against IFN-γ restriction of bacterial growth (**Fig. 5A**). We pretreated *CYLD* KO cells, or a non-target control cell line, with IFN-γ for 24 h before infecting them with *L. monocytogenes* and examined growth by plating for colony-forming units (CFU) after 6 hpi. No differences in growth were observed between the non-target and *CYLD* KO cell lines, suggesting that CYLD does not protect *L. monocytogenes* from bacterial clearance in epithelial cells (**Fig. 5B**). We also examined M1-linked ubiquitination and p62 localization of *L. monocytogenes* in IFN-γ-treated *CYLD* KO cells by microscopy. We did not observe changes in M1-linked ubiquitin or p62 localization to *Lm*Dasher in the *CYLD* KO cells (**Fig. 5C** - **F**), Therefore, despite InlC recruiting CYLD to ubiquitinated *L. monocytogenes*, CYLD does not deubiquitinate intracellular bacteria or regulate the recruitment of autophagic markers to the pathogen, highlighting a potential cell-type specific role for CYLD beyond limiting bacterial clearance.

**Figure 5.**
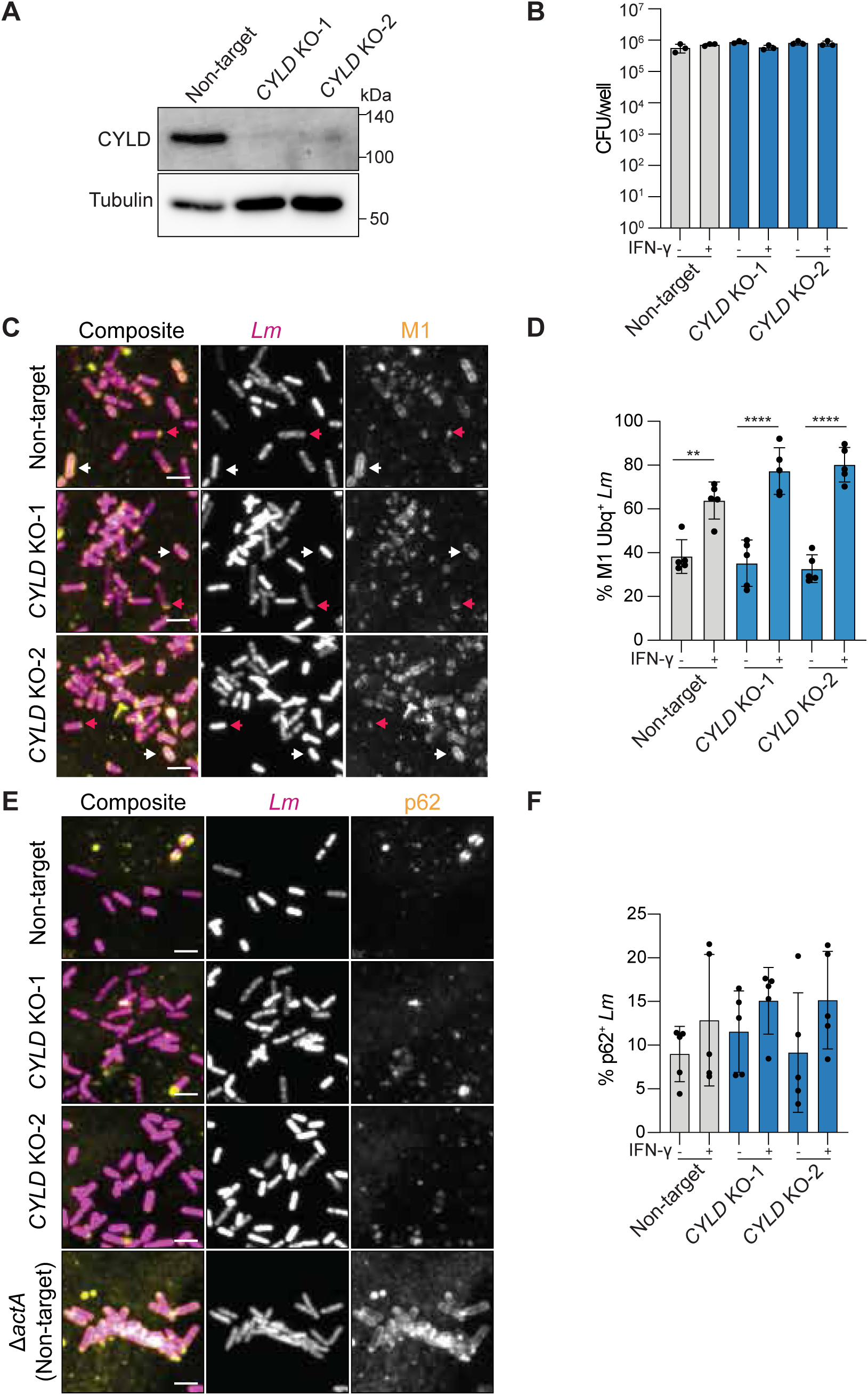
CYLD does not impact bacterial growth, ubiquitination, or p62 co-localization in A549 epithelial cells. **(A)** Western blot analysis of CYLD from A549 *CYLD* KO or non-target control cells. (**B**) Enumerated colony-forming units (CFU) per well of *L. monocytogenes* from infected *CYLD* KO cells with or without IFN-γ. (**C** - **F**) Immunofluorescence detection and percentage of *Lm*Dasher co-localized with M1-linked ubiquitin (**C** and **E**) or p62 (**E** and **F**) at 5 hpi in A549 *CYLD* KO cells with and without IFN-γ. *Lm*Dasher (magenta); anti-M1-linked ubiquitin (**C**) and anti-p62 (**E)** (yellow). Arrows indicate examples of polar (red) and uniform (white) M1-linked ubiquitin. Non-target cells infected with Δ*actA Lm*Dasher was used as a control for p62 localization. All data are representative of two (**C** and **D**) and three (**E** and **F**) independent experiments and plotted as the percentage of indicated bacteria per FOV displayed as mean ± SD; >240 bacteria were counted per FOV in each experiment. Scale bar, 2 μm. Significance determined by Ordinary one-way ANOVA using Tukey’s multiple comparisons post-hoc test \*\**P*<0.01, ****P<0.0001.

### InlC co-opts CYLD to protect against IFN-γ restriction of cell-to-cell spread

In epithelial cells, InlC promotes *L. monocytogenes* cell-to-cell spread (34–36), and we next asked if InlC and CYLD cooperate to further enhance this process. To test CYLD’s role in spread, we performed an infectious focus assay at 7 hpi using *Lm*Dasher at a very low multiplicity of infection (MOI). Assays were performed with *CYLD* KO and non-target control cell lines pretreated with PBS or IFN-γ for 24 h before infection. IFN-γ treatment caused a modest reduction in cell-to-cell spread in the non-target, control cells (**Fig. 6A**), suggesting it may restrict dissemination. Loss of *CYLD* expression also caused a slight reduction in cell-to-cell spread in both *CYLD* KO lines compared to non-target controls (**Fig. 6A**). Interestingly, loss of *CYLD* expression dramatically sensitized *L. monocytogenes* to IFN-γ, leading to a significantly larger reduction in cell-to-cell spread in both *CYLD* KO cell lines (**Fig. 6A**). Therefore, while CYLD promotes spread in the absence of IFN-γ treatment, it also protects *L. monocytogenes* from IFN-γ restriction of cell-to-cell spread.

**Figure 6.**
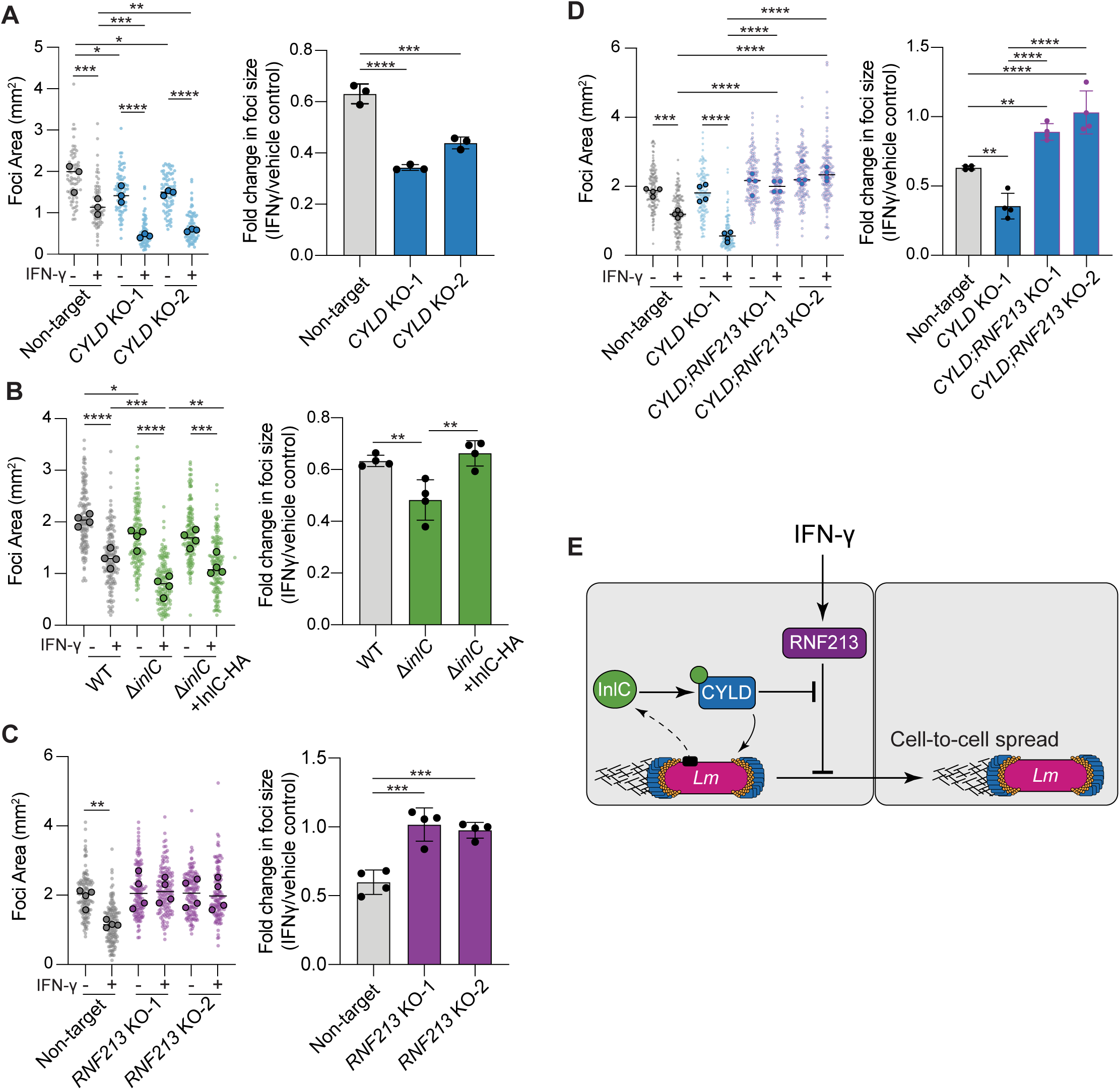
CYLD protects against IFN-γ-mediated, RNF213-dependent restriction of cell-to-cell spread. (**A** - **D**) Area of infectious foci from IFN-γ pretreated (**A**) A549 *CYLD* knockout cells infected with *Lm*Dasher, (**B**) A549 cells infected with wild type, *ΔinlC*, or *ΔinlC* + HA-InlC complemented strain of *L. monocytogenes,* (**C**) A549 *RNF213* knockout cells infected with *Lm*Dasher, or (**D**) from A549 *CYLD* and *RNF213* double knockout cells infected with *Lm*Dasher for 7 h. The left plots display medians from 3 – 4 independent experiments (dark circles) superimposed on the raw data (transparent circles) and were used to calculate the median (black line) and *P*-values. The right plot displays the fold change of median foci size between IFN-γ treatment and vehicle control. Significance determined using Ordinary one-way ANOVA using uncorrected Fisher’s least significance difference (LSD) post-hoc test \**P*<0.05, \*\**P*<0.001, \*\*\**P*<0.0001, \*\*\*\**P*<0.00001. (E) Schematic of InlC and CYLD suppression of IFN-γ-mediated RNF213 restriction of cell-to-cell spread.

We next wanted to determine whether InlC similarly protected the bacteria against IFN-γ-mediated restriction of spread. We performed an infectious focus assay in IFN-γ-pretreated A549 cells infected with either wild-type, Δ*inlC*, or the Δ*inlC* + HA-InlC complement strain of *L. monocytogenes.* Similar to prior studies, deletion of *inlC* resulted in a slight reduction in cell-to-cell spread compared to wild-type *L. monocytogenes* in vehicle-treated cells (**Fig. 6B**). As observed with the *CYLD* KO cells, IFN-γ induced a significantly greater reduction in spread by the Δ*inlC* strain compared to wild-type *L. monocytogenes*, which was especially apparent when comparing the fold change in spread between treated and untreated samples (**Fig. 6B**). Furthermore, this IFN-γ exacerbated defect was restored when HA-InlC was complemented *in trans*, though this expression was insufficient to fully complement the loss of *inlC* in vehicle-treated cells (**Fig. 6B**). Collectively, these data suggest that InlC and CYLD protect *L. monocytogenes* from IFN-γ restriction of cell-to-cell spread.

*L. monocytogenes* spreads from cell to cell via a polar actin tail (2), a site at which CYLD is also localized. IFN-γ treatment caused a significant reduction in actin tail frequencies in non-target cells (**Fig. S3**), which prompted us to asked whether CYLD primarily acted by protecting actin-based motility from IFN-γ-mediated restriction. Instead, we found that loss of *CYLD* expression did not consistently impact tails in untreated conditions, and it did not exacerbate the INF-γ-induced defect (**Fig. S3**). Thus, CYLD’s protection against IFN-γ and its restriction of cell-to-cell spread cannot be attributed solely to inhibition of actin tail formation.

RNF213 is an IFN-γ inducible gene that restricts bacterial growth in macrophages (26, 37, 38), but we did not detect a similar IFN-γ-induced growth defect in epithelial cells (**Fig. S4**). Since we found that RNF213 regulates IFN-γ dependent M1-linked ubiquitination and CYLD re-localization to *L. monocytogenes*, we examined whether RNF213 also regulated IFN-γ restriction of cell-to-cell spread by performing an infectious focus assay in vehicle and IFN-γ-treated *RNF213* KO cells. In the absence of IFN-γ, the loss of *RNF213* expression had no impact on cell-to-cell spread (**Fig. 6C**), indicating it does not promote spread in untreated conditions. However, upon IFN-γ treatment, loss of *RNF213* expression led to normal levels of spread, indicating it is necessary for IFN-γ restriction of spread. Given the opposing roles of CYLD and RNF213 during spread, we then asked if CYLD counteracted the RNF213-mediated restriction induced by IFN-γ. To investigate this idea, we generated A549 *CYLD* and *RNF213* double-KO cells (*CYLD*;*RNF213* KO-1 and *CYLD*;*RNF213* KO-2) (**Fig. S5**) and examined whether loss of both CYLD and RNF213 abolished IFN-γ restriction of spread. Treating *CYLD* KO lines with IFN-γ continued to show a significant reduction in cell-to-cell spread compared to non-target controls (**Fig. 6D**). However, no difference in cell-to-cell was observed when comparing vehicle and IFN-γ-treated *CYLD*/*RNF213* double-KO cell lines (**Fig. 6D**). Therefore, these results demonstrate *L. monocytogenes* co-opts CYLD to counteract IFN-γ-mediated, RNF213-dependent restriction of cell-to-cell spread (**Fig. 6E**).

## DISCUSSION

*L. monocytogenes* recruits host proteins to its surface to evade immune defenses and move within the host (2, 6). Defining the host-pathogen interactome at the bacterial surface, and understanding how these interactions vary across host cell types is critical for advancing our understanding of pathogenesis. Here, we applied the proximity-dependent biotin ligase split-TurboID to profile host proteins recruited to the *L. monocytogenes* surface during intracellular infection. We discovered that the host deubiquitinase CYLD is recruited to *L. monocytogenes*, and that in nonphagocytic cells, it counteracts IFN-γ-mediated, RNF213-dependent restriction of cell-to-cell spread. Furthermore, we demonstrate that CYLD recruitment to the bacterial surface depends on the secreted effector InlC, which itself promotes cell-to-cell spread, thereby linking effector-mediated host protein recruitment to intercellular dissemination. Altogether, these data underscore the value of pathogen surface profiling in uncovering previously unappreciated roles for host factors during the *L. monocytogenes* infectious life cycle.

Previous work established a critical role for CYLD during infection with various pathogens like *L. monocytogenes*, *Staphylococcus aureus*, and *Haemophilus influenzae* (39–41). Indeed, loss of *CYLD* expression in macrophages or *in vivo* led to enhanced killing of *L. monocytogenes* and *S. aureus*, suggesting pathogens co-opt CYLD to promote infection in phagocytes (21, 39). CYLD also deubiquitinates STAT3 to reduce fibrin production, reduce neutrophil recruitment, and promote *L. monocytogenes* dissemination in the liver (21). In our study, loss of *CYLD* expression in epithelial cells did not restrict *L. monocytogenes* burdens, even though CYLD is known to regulate innate immune signaling in these cells (21, 40). Instead, our work shows *L. monocytogenes* relies on CYLD to promote cell-to-cell spread. In addition to its canonical role as a negative regulator of innate immune signaling, CYLD regulates cell cycle progression and promotes gap junction formation through the deubiquitination of plakoglobin in a variety of epithelial and endothelial cells (42, 43). CYLD also controls ciliogenesis in numerous cell types through deubiquitination of centrosomal proteins such as Cep70 and negative regulation of the histone deacetylase HDAC6 (44). How CYLD is regulated to carry out these diverse functions across various cell types remains poorly understood, but our data suggest *L. monocytogenes* leverages CYLD’s cell-type-specific roles to promote infection. Future work is needed to examine how this specificity is achieved and how pathogens like *L. monocytogenes* exploit these differences during systemic disease.

We demonstrate that InlC binds to and re-localizes CYLD to M1-linked ubiquitin at the poles of the bacteria, but how InlC regulates CYLD’s localization remains an open question. InlC and CYLD did not co-localize at sites of CYLD enrichment around the bacteria, suggesting InlC’s interaction with CYLD may be transient or mediated by a minor fraction of the secreted pool. InlC was also sufficient to enable CYLD localization to other ubiquitinated bacterial species, suggesting InlC alone alters CYLD activity. InlC has been shown to bind several host targets, including IKKα to suppress NF-κB signaling, and the cytoskeleton adapter TUBA and exocyst complex members to promote cell-to-cell spread (33–35). CYLD’s substrate specificity, activity, and subcellular localization are regulated by IKKε- and IKKβ-mediated phosphorylation (45–47), which are kinases that also cooperate with InlC’s binding partner IKKα (33, 29). Thus, further studies are needed to examine what impact InlC has on the post-translational modifications of CYLD and if this contributes to CYLD localization and function.

RNF213 localizes to *L. monocytogenes* (37) and our study showed that it was responsible for M1-linked ubiquitination of the bacterial poles. This work expands the range of pathogens ubiquitinated by RNF213, as prior work showed that RNF213 directly ubiquitinates the lipid A moiety of lipopolysaccharide on *Salmonella enterica* serovar Typhimurium (26). Other E3 ligases, such as RNF166, also co-localize with *L. monocytogenes*, but they enhance autophagic clearance by ubiquitination of p62 and not a bacterial substrate (9). It remains to be determined whether RNF213 targets bacterial proteins or other non-proteinaceous substrates on the surface of Gram-positive bacteria, or other host proteins recruited to the bacterial surface.

We found that RNF213 and CYLD play opposing roles in regulating cell-to-cell spread, but loss of *CYLD* expression had only a minimal effect on ubiquitination of *L. monocytogenes*. How then does CYLD limit RNF213-mediated restriction of cell-to-cell spread? Innate immune signaling (e.g., via IFN-γ) may trigger ubiquitination of substrates that CYLD must derepress to promote dissemination. While CYLD does not protect against IFN-γ-mediated restriction of actin tail formation, CYLD could deubiquitinate specific cytoskeletal regulators at the bacterial surface, thereby affecting the speed, direction, or stability of actin tails during spread. Alternatively, CYLD recruitment to the pathogen surface may precede its activity elsewhere in the cell that contributes to de-repressing factors that restrict the spread. Exploring how infection shapes the ubiquitin landscape and the interplay between CYLD and RNF213 activities will be necessary to understand how specific ubiquitinated factors regulate bacterial dissemination during infection.

Overall, our work demonstrates that surface display of promiscuous biotin ligases can be used to profile the bacterial surface during intracellular infection, revealing unexpected contributions of mammalian proteins to the pathogenesis of *L. monocytogenes*. While our study focused on CYLD, the expanded list of interacting host factors identified in our screen will motivate future work into their roles during infection. With this tool in hand, researchers will also be able to probe subcellular-specific changes to the interactome by substituting the cytosol-targeted CTurboID with one that is recruited to host cell compartments or membranes. Dynamic or cell-type specific changes to the pathogen surface could also be profiled using different time points or cellular backgrounds. While our split-TurboID approach successfully captured novel, pathogen-associated regulators of cell-to-cell spread, we note that very few *L. monocytogenes* proteins were identified (**Table S2**). Future iterations of this tool may address this by modifying the lysis conditions or exchanging the ligase used to improve pathogen labeling. Such advances may also inform how similar approaches could be applied to other pathogens. Given the growing interest in directly probing the pathogen surface during infection, continued optimization and application of proximity-dependent profiling approaches offer a powerful strategy to reveal needed insights into the diversity of host-pathogen interactions within and across distinct cellular environments.

## METHODS

### Cell lines

A549 (human lung epithelial cells; RRID:CVCL_A549), HEK293T (human embryonic kidney, RRID:CVCL_0063), and Vero (monkey kidney epithelial, RRID:CVCL_0059) cells were obtained from the University of California, Berkeley Cell Culture Facility (Berkeley, CA USA). Cell lines were cultured in high-glucose Dulbecco’s Modified Eagle Medium (DMEM, 11965118, Gibco ThermoFisher Scientific) supplemented with either 10% (A549 and HEK293T cells) or 5% (Vero cells) fetal bovine serum (FBS, Gibco ThermoFisher Scientific #A5670701) at 37°C with 5% CO_2_.

### Bacterial strains

All bacterial strains and plasmids used in this study are listed in Table S3. *Listeria monocytogenes* serotype 1/2a (serotype 10403s) and derived mutants were grown on brain heart infusion (BHI) broth or agar at 30°C. *Escherichia coli* DH5α and SM10 (a kind gift from Daniel Portnoy, University of California, Berkeley) were grown on Luria-Bertani (LB) broth or agar at 37°C. *Rickettsia parkeri* GFP (*Rp*GFP) and the *ompB*^STOP^::Tn were generated as previously described (30, 48). All *R. parkeri* strains were grown at 33°C in Vero cells with DMEM supplemented with 2% FBS and extracted from host cells by mechanical disruption as previously described (48, 49).

### Generation of plasmids

To generate NTurboID surface display constructs, gene fragments were synthesized (TWIST Biosciences) containing the *actA* promoter followed by the signal peptide sequences of either InlA, InlF, or ActA, NTurboID-HA codon optimized for *L. monocytogenes*, and the cell wall or membrane anchors of the indicated surface proteins. These fragments were cloned into the *L. monocytogenes* integration pPL2x (17). To generate the surface display construct with iActA linker (pPL2x_ActApr-NTurboID-HA-ActAi-ActAtm) the disordered region of *L. ivanovii* ActA was amplified from the genome of *L. ivanovii* FSL C2-010 (a kind gift from Darren Higgins, Harvard Medical School, Boston, MA) and inserted between NTurboID-HA and the membrane anchor of ActA in pPL2x. pPL2X_ActApr_DasherGFP was generated by amplifying *Dasher*GFP and the associated upstream *actA* promoter, from *Lm*Dasher and inserted into pPL2x. The pPL2x_InlCpr_InlC-HA complement vector was generated by amplifying InlC with its endogenous promoter from the *L. monocytogenes* genome with primers containing a sequence for an HA-tag and inserted into pPL2x.

For RDA_550 knockout vectors, synthesized gRNA (Millipore Sigma) containing two tandem guides per construct, chosen from the Humagne C+D gRNA library (50) were inserted into BsmBI-digested pRDA_550 using Golden Gate cloning protocols (51). Guide sequences are listed in Table S4. Non-target sequences were cloned into pRDA_550 as previously published (52).

The lentiviral expression vector FUW2IB_CTurboID_V5_NES was generated by synthesizing a gene fragment containing CTurboID fused to a V5 tag into FUW2IB. This construct was modified by PCR-mediated addition of a nuclear export signal. To express HaloTag and HaloTag-CYLD in mammalian cells, HaloTag and HaloTag-CYLD were amplified by PCR and inserted into FUW2IB. Finally, the pCLIP2B_HA_InlC oncoretroviral expression vector was constructed by inserting a synthesized gene fragment (TWIST Biosciences) containing human codon-optimized InlC with an N-terminal HA-tag into pCLIP2B.

### Generation of *Lm*Dasher, LmNTurboID, InlC complemented strains of *L. monocytogenes*

All pPL2x-derived constructs were integrated into the PSA prophage integration site at the 3’ of the arginine tRNA gene through conjugation, as previously described (17). In brief, all generated pPL2x constructs were transformed into SM10 *E. coli*. These donor strains were patched on BHI agar with the appropriate *L. monocytogenes* parental strain and incubated for 24 h at 37°C. Conjugative patches were then restreaked for individual colonies on BHI agar supplemented with 200 µg/mL streptomycin and 2 µg/mL tetracycline and incubated at 37°C for 36 – 48 h until visible colonies appeared. Individual colonies were streaked onto antibiotic-supplemented BHI agar and incubated at 37 °C for 18 h to confirm resistance. Strains were maintained in antibiotics until saved as glycerol stocks. Functional validation of NTurboID surface display and *Lm*Dasher were confirmed by confocal microscopy as described below. Validation of Δ*inlC* + InlC-HA complement strain was done by immunofluorescence and western blotting for InlC-HA.

### Generation of A549 cells stably expressing CTurbo-V5-NES, HaloTag, HaloTag-CYLD, or HA-InlC

All stable cell lines were generated using lentiviral or oncoretroviral transduction as previously described (48). To generate A549-CTurboID, A549-HaloTag, and A549-HaloTag-CYLD cell lines, viral particles were packaged by transfecting 2.5 x 10^5^ cells/mL (2mL, 6-well plate) of HEK293T seeded the day before with 200ng each of pMDL/RRE, pRSV/REV, and pCMV-VSVG packaging plasmids along with 400ng of either CTurbo-V5-NES_FUW2IB or HaloTag-CYLD_FUW2IB using calcium phosphate. 24 hours after transfection, the medium was discarded and replaced with 1.5mL DMEM supplemented with 10% FBS, and the cells were incubated for an additional 24 hours. Lentiviral media were harvested and filtered through a 0.45 µM filter before being added to A549 cells seeded at 0.9 × 10^^5^ cells/mL (2mL, 6-well plate) the day before. After 24 h, 1mL of DMEM supplemented with 10% FBS was added to transduced cells and incubated for an additional 24 h. Lentiviral media was then discarded and replaced with 2mL DMEM supplemented with 10% FBS and 12 µg/mL blasticidin (R21001; ThermoFisher Scientific) to select for transduced cells. A549-HaloTag-CYLD that also stably expresses HA-InlC or pCLIP2B empty vector control was generated as described above, except 200ng of pAmpho and pCMV-VSVG were used as packing vectors, and 1.5 µg/mL puromycin (p8833; Sigma-Aldrich) was used for selection.

### Generation A549 RNF213, HOIP (RNF31), CYLD and RNF213/CYLD double knockout cell lines

All knockout cell lines were generated using lentiviral transduction of CRISPR Cas12a (51). Viral particles were packaged by transfecting 0.66 x 10^5^ cells/mL (0.5mL, 24-well plate) of HEK293T seeded the day before with 140ng each of MDL/RRE, pRSV/REV, and pCMV-VSVG packaging plasmids along with 280ng of RDA_550 EnAsCas12a guide containing vector using the TransIT-LT1 transfection reagent (Mirus Bio, #2304) and centrifuged at 1000 × *g* for 30 minutes at room temperature. 6 h after transfection, the media was discarded and replaced with 1mL DMEM supplemented with 10% FBS and 1% bovine serum albumin (BSA), and incubated for 36 h. After incubation, viral media was removed, filtered through a 0.45 µM filter, and 0.2mL of filtered viral media was then added to A549 cells seeded at 0.9 x 10^4^ cells/mL (2mL, 6-well plate), seeded the day before. Transduction media was further supplemented with 8 µg/mL polybrene (TR-1003-G, EMD Millipore), centrifuged at 1,000 × *g* for 30 minutes, and incubated for 4 - 6 h. After initial incubation, cells were washed with PBS and were recovered in 2mL DMEM supplemented with 10% FBS for 24 h before adding 1.5 µg/mL puromycin to select for transduced cells.

To confirm successful knockout of the indicated genes, puromycin-selected cells were seeded at 2.5 × 10^5^ cells/well (0.5mL, 24-well plate) and incubated for 5 days, as described for bacterial infections below. After 5 days, cells were washed with 1X PBS and harvested by scraping in RIPA buffer (50mM Tris, pH = 7.4, 150mM NaCl, 1% NP-40, 0.5% sodium deoxycholate, 0.1% sodium dodecyl sulfate) supplemented with protease inhibitor cocktail (K1008, APEXBio). Samples were incubated while rotating at 4°C for 15 min, centrifuging at 10000 × *g* at 4°C for 15 min, and the clarified lysate was collected for analysis. Protein concentrations were measured and normalized using the Pierce BCA Assay (23227; ThermoFisher Scientific) and analyzed by western blot using rabbit anti-HOIP (ab46322; abcam), rabbit anti-RNF213 (HPA026790; Millipore Sigma), rabbit anti-CYLD (8462S; Cell Signaling Technology), and mouse anti-Tubulin (T6199, Sigma-Aldrich).

### Bacterial infection of cultured cells

The indicated strain of *L. monocytogenes* was streaked onto BHI and incubated at 30°C for 20 h. The following day, 5 colonies of *L. monocytogenes* were used to inoculate 2.5mL BHI broth and incubated as a stationary culture at 30°C for 20 h. After overnight growth, 1mL of culture was centrifuged at 16,200 × *g* for 1 min and resuspended in PBS. A549 cells seeded at 2.5 x 10^5^ cells/well (0.5 mL, 24-well plate) or 2.8 x 10^5^ cells/mL (2mL, 6-well plate) 5 days prior were infected at the indicated multiplicity of infection (MOI), centrifuged at 30 × *g* for 5 minutes, and incubated for 1 h to allow for bacterial invasion. After initial incubation, the media was discarded, cells were washed twice with 1X PBS, and given fresh DMEM supplemented with 10% FBS and 10 μg/mL gentamicin. Infected cells were further incubated for 4 – 6 h as indicated and processed for the indicated downstream assay.

### Examining surface display of NTurboID on *L. monocytogenes*

NTurboID-HA was detected in *Lm*NTurboID-infected A549 cells using a two-step permeabilization and immunofluorescence protocol. The first step permeabilizes host cells only, which means only bacterial surface proteins exposed to the host cell cytosol are detected. The second step permeabilizes bacteria to detect proteins within the bacteria. The second step was used as a control to ensure any negative result from staining the surface-exposed NTurboID was not due to NTurboID-HA getting trapped within the bacteria. A549 cells seeded at 2.5 x 10^5^ cells/well (0.5mL, 24-well plate) on glass cover slips were infected with *Lm*TurboID at an MOI of 1. After incubation, cells were washed twice with 1X PBS and fixed with 4% paraformaldehyde (PFA) in PBS for 0.5 – 1 h. Fixed cells were washed three times with 1X PBS, incubated in 0.1 M glycine/PBS for 10 min, washed again with 1X PBS, and then permeabilized with 0.5% Triton-X 100 in 1X PBS for 5 minutes. Samples were then washed three times with PBS and blocked in blocking buffer (2% BSA and 10% normal goat serum in 1X PBS) for 30 min, followed by incubation with mouse anti-HA (901533; Biolegend) and rabbit Listeria O Antisera (22301; BD Difco) diluted at 1/1000 and 1/200, respectively, in blocking buffer for 1 h. After primary incubation, cells were washed three times with 1X PBS and then incubated in goat anti-rabbit secondary antibody conjugated with Alexa Fluor 488 (A-11008; ThermoFisher Scientific) and goat anti-mouse secondary antibody conjugated with Alexa Fluor 568 (A-11004; ThermoFisher Scientific) both diluted 1/200 in blocking buffer for 1 h. After secondary incubation, coverslips were washed three times with 1X PBS followed by crosslinking antibodies 4 % PFA/PBS for 5 minutes. Coverslips were washed again with PBS before quenching the additional PFA by incubating the coverslips again in 0.1M glycine/PBS for 5 min. Coverslips were washed three times in 1X PBS and then incubated in lysozyme reaction buffer (0.8X PBS, 50mM glucose, 5mM EDTA) supplemented with 5mg/mL lysozyme (L6876; Sigma Aldrich) and 0.1% Triton-X 100 for 30 minutes at 37°C to permeabilize the bacteria. Coverslips were washed three times before being incubated with mouse anti-HA diluted 1/1000 in blocking buffer for 3 hours at 37°C. Coverslips were washed three times before being incubated in goat anti-mouse secondary antibody conjugated with Alexa Fluor 405 (A48255; ThermoFisher Scientific) diluted 1/200 in blocking buffer for 1 hour at room temperature. After secondary incubation, cells were washed three times with 1X PBS before being mounted on glass slides with ProLong Gold Antifade Mountant (P36934; Invitrogen). Stained slides were then imaged with the Olympus IXplore Spin microscopy system, Yokogawa CSU-W1 spinning disk unit, and an ORCA-Flash4.0 sCMOS 537 camera using a 100x UPlanSApo (1.35 NA) objective.

### Biotin labeling with split-TurboID

To examine biotin labeling by confocal microscopy, A549-CTurboID cells seeded on glass cover slips were infected with *Lm*TurboID at an MOI of 5. After the media was discarded and the cells were washed, the cells were supplemented and incubated for 2 h. The cells were then treated with 50 µM biotin or 0.1% DMSO vehicle control and incubated an additional 2 h. After incubation, cells were washed twice with 1X PBS and fixed with 4% PFA/PBS for 0.5 – 1 h. Fixed cells were washed three times with 1X PBS, incubated in 0.1 M glycine/PBS for 10 min at room temperature, washed again with 1X PBS, and then permeabilized with 50 µg/mL digitonin in 1X PBS for 10 min. Samples were washed three times with PBS and blocked in blocking buffer (2% BSA in 1X PBS) for 30 min. Samples were then incubated with rabbit Listeria O Antisera at 1/200 dilution in blocking buffer for 1 h. After primary incubation, cells were washed three times with 1X PBS and then incubated in goat anti-rabbit secondary antibody conjugated with Alexa Fluor 488, and Neutravidin conjugated with Rhodamine Red (A6378; ThermoFisher Scientific) diluted 1/200 and 1/1000, respectively, in blocking buffer for 1 h. After secondary incubation, cells were washed three times with 1X PBS before being mounted on glass slides with ProLong Gold Antifade Mountant and imaged with the Olympus IXplore Spin microscopy system using a 100x objective.

To examine biotin labeling by western blot, A549-CTurboID cells were infected with *Lm*TurboID as described, except 50 µM biotin or 0.1% DMSO vehicle control was supplemented in the media immediately after PBS washes, following the 1 h incubation for invasion, and incubated for an additional 5 hours. After biotin incubation (6 hpi), cells were washed and harvested by scraping in RIPA buffer supplemented with protease inhibitor cocktail (K1008, APEXBio). Samples were incubated while rotating at 4°C for 15 min, centrifuging at 10000 × *g* at 4°C for 15 min, and the clarified lysate was collected for analysis. Protein concentrations were determined and normalized using the Pierce BCA Assay (23227; ThermoFisher Scientific), and analyzed by western blot using streptactin-HRP (1610381; BioRad), mouse anti-HA (901533; Biolegend), mouse anti-V5 (R960-25; ThermoFisher Scientific), and mouse anti-GAPDH (2118T: Cell Signaling Technology).

### Streptavidin enrichment and identification of host proteins at the bacterial surface

A549-CTurboID cells were seeded at 2.8 x 10^5^ cells/mL (2mL, 20 x 6-well plates) and half the plates were infected with wild-type or ΔactA *Lm*NTurboID at an MOI of 5. After invasion, the cells were treated with 50 µM biotin and incubated for 5 h. The cells were washed twice with cold 1X PBS and harvested by scraping in RIPA buffer. Lysates from 2.5 plates were pooled as individual biological replicates and processed as done above for western blotting. 90 μL of streptavidin magnetic bead (88817; ThermoScientific) were equilibrated by washing twice with RIPA buffer for 2 minutes while rotating at 4°C. Beads were then incubated in 2mL of 3.5mg/mL of lysate overnight at 4°C while rotating. Beads were then washed twice with 1mL of RIPA buffer for 2 minutes, once with 1M KCl for 2 minutes, once with 0.1M sodium carbonate for 1 minute, once with 2M Urea (in 10mM Tris pH = 8.0) for 1 minute, followed by two additional washes in PBS for 2 minutes. All washes were performed at 4 °C while rotating. After the last wash, beads were processed for mass spectrometry.

Protein-bound magnetic bead samples were mixed 1:1 by volume with 1% SDC in 100mM TEAB, 40mM CAA, and 10mM TCEP, and reduced and alkylated at 70 °C for 15 min. Proteins were digested overnight with 1 µg trypsin/Lys-C mix in 100mM TEAB at 37 °C in a shaking incubator at 115 RPM. The following day, an additional 1 µg dose of trypsin/LysC mix was added in 100mM TEAB and the digestion continued at 37 °C for 4 hours. Peptide digests were purified with stage tips as described in Rappsilber *et al*. in 2007 (53). Peptides were then dried in a speed-vac concentrator at 50 °C and resuspended in 0.2% (v/v) formic acid in MS-grade water for LC-MS analysis. LC-MS/MS data were collected on an Orbitrap Eclipse mass spectrometer coupled with a Vanquish Neo nanoLC, a FAIMS Pro Interface, and an Easy Spray ESI source, all by Thermo Fisher Scientific (Waltham, MA, USA). NanoLC separation utilized an EasySpray ES902 column (75 µm x 25 cm, 100 Ȧ) from Thermo Fisher Scientific. An injection volume of 5 µL was used for peptide extracts. The mobile phase consisted of 0.1% (v/v) formic acid in water (solution A) and 0.1% (v/v) formic acid in 80% (v/v) acetonitrile (solution B), separating at 300 nL/min and maintaining a column temperature of 50 °C. Initial column conditioning was performed with 3% solution B, followed by a linear gradient increasing to 25% B over 45 minutes, then to 40% B over 15 minutes, and finally to 95% B over 5 minutes. Remaining peptides on the C18 resin were eluted at 95% B for 6 minutes.

In positive ion mode, the ion source temperature was set at 305 °C with a voltage of 2000 V, and ionized peptides passed through the FAIMS Pro unit at compensation voltage -50 V. MS1 spectra were acquired with a resolution of 120,000 and a mass range of 350-2000 m/z. Standard automatic gain control settings and automatic maximum injection times were used. For MS2 acquisition, the mass spectrometer was operated in DIA mode with a resolution of 30,000, gathering spectra across a precursor mass range of m/z 375-1200.

Isolation windows of m/z 25 were used, with 0.5 m/z overlaps, a custom AGC target of 1000 units, and 30% normalized CID collision energy.

The DIA-NN 1.8 software platform (54) was utilized to analyze proteome data using the default settings for the software using standard settings. Library generation and library-free searches were conducted using the *Listeria monocytogenes* serotype 1/2a FASTA database (UP000001288) and Human (Uniprot). The resulting quantitative protein data were exported in .tsv format and analyzed according to the methods described by Schulte *et al*. 2019 (55). Normalized data were used to calculate log_2_fold changes and adjusted P-values using a two-stage setup method of Benjamini, Krieger, and Yekutieli.

### Examining HaloTag-CYLD, ubiquitin, HA-tagged InlC, and actin tail localization by microscopy

Localization of ubiquitin was examined by seeding A549 cells, or the indicated knockout cell line, at 2.5 x 10^5^ cells/well (0.5mL, 24-well plate) on glass coverslips and infected with *Lm*Dasher at an MOI of 0.25. Where indicated, cells were treated with recombinant 25 ng/mL IFN-γ (300-02; ThermoFisher Scientific) or PBS vehicle control 24 h before infection. Infected cells were fixed with 4% paraformaldehyde at 5 hpi (4 h post invasion media change). Fixed cells were washed three times with 1X PBS, incubated in 0.1 M glycine/PBS for 10 min at room temperature, washed again with 1X PBS, and then permeabilized in 0.5% Triton-X 100 in 1X PBS for 5 minutes. Samples were washed three times with PBS and blocked in blocking buffer for 30 min. Samples were incubated with mouse anti-ubiquitin FK1 (04-262; Millipore Sigma) and rabbit anti-M1-linked ubiquitin (ZRB2114; Millipore Sigma), both diluted 1/200 in blocking buffer for 1 h. After primary incubation, cells were washed three times with 1X PBS and then incubated in goat anti-rabbit secondary antibody conjugated with Alexa Fluor 405 and goat anti-mouse secondary antibody conjugated with Alexa Fluor 568 or Alexa Fluor 647, all diluted 1/200 in blocking buffer for 1 h. After secondary incubation, samples were washed three times with 1X PBS before being mounted on glass slides with ProLong Gold Antifade Mountant (P36934; Invitrogen). Stained slides were then imaged with the Olympus IXplore Spin microscopy system using a 100x objective.

To examine the localization of HaloTag-CYLD and HA-InlC, A549-HaloTag-CYLD or A549-HaloTag-CYLD + HA-InlC cells were infected with either *Lm*Dasher or *Lm*BFP on glass coverslips as done with ubiquitin staining. For experiments examining HaloTag-CYLD around *R. parkeri,* cells were infected as done with *L. monocytogenes* at an MOI of 1 - 5. At 4.5 hpi, cells were supplemented with Janelia Fluor 635 HaloTag Ligand HT1050; Promega) to image HaloTag-CYLD and incubated for an additional 30 min before fixation. Samples were further stained and mounted on glass slides as described above using mouse anti-HA (901533; Biolegend) diluted in 1/1000 in blocking buffer and goat anti-mouse secondary antibody conjugated to Alexa Fluor 647 diluted 1/200 in blocking buffer when examining HA localization. Stained slides were then imaged with the Olympus IXplore Spin microscopy system using a 100x objective.

Finally, to quantify actin tails in *CYLD* KO cells, non-target or *CYLD* KO cells were seeded at 2.5 x 10^5^ cells/well (0.5mL, 24-well plate) on glass coverslips and infected with *Lm*GFP at an MOI of 0.25. Where indicated, cells were treated with recombinant 25 ng/mL IFN-γ or PBS vehicle control 24 h before infection. Infected cells were fixed with 4% PFA/PBS at 5 hpi (4 h post invasion media change). Coverslips were stained and mounted on glass slides as described above, using phalloidin conjugated to iFluor 405 (ab176752; abcam) diluted 1/100 in blocking buffer to stain for actin. Stained slides were then imaged with the Olympus IXplore Spin microscopy system using a 100x objective.

Localization of ubiquitin, HaloTag-CYLD, and actin tails are expressed as a percentage of the total of all counted *L. monocytogenes* per FOV within an independent experiment. All data were quantified manually using ImageJ.

### Infectious focus assay to measure bacterial spread

The indicated A549 knockout cell line was seeded at 2.5 x 10^5^ cells/well (0.5mL, 24-well plate) on glass coverslips and infected with *Lm*Dasher at an MOI of 0.005 as described above. Where indicated, cells were treated with recombinant IFN-γ or PBS vehicle control 24 h before infection. Infected cells were fixed 7 hpi with 4% PFA/PBS and coverslips were stained as described for examining protein localization using mouse anti-β-catenin (2677: Cell Signaling Technology) diluted at 1/200 in blocking buffer along with Hochest (H3570; ThermoFisher Scientific) diluted 1/1000, phalloidin conjugated to Alexa Fluor (A22287; Fisher Scientific) diluted 1/200, and goat anti-mouse secondary antibody conjugated to Alexa Fluor 568 diluted 1/200. Stained slides were then imaged with the Olympus IXplore Spin microscopy system using a 60x UPlanSApo (1.30 NA) objective.

To measure the area of infectious foci, 24 - 36 individual foci were imaged for each replicate (n = 3 - 4). A binary gray-scale max projection of the bacteria-containing channel was generated for each foci imaged and used to measure the area covered by segmented bacteria using AlphaShapes with an α = 4.5 as previously described (56).

### HaloTag pulldown

A549, A549-HaloTag, or A549-HaloTag-CYLD cells were seeded at 2.8 x 10^5^ cells/mL (2mL, 6-well plate) and infected with Δ*inlC* + InlC-HA *L. monocytogenes* at an MOI of 5. At 6 hpi, cells were washed with 1X PBS and harvested by scraping in immunoprecipitation lysis buffer (50mM Tris-HCl pH=7.4, 150 mM NaCl, 1 mM EDTA, 1% Triton X-100) supplemented with protease inhibitor and incubated for 15 minutes while rotating at 4°C. Lysates were then centrifuged at 11300 × *g* for 15 minutes at 4°C. 1 mg of clarified lysate was added to 25uL of equilibrated ChromoTek Halo-Trap magnetic agarose beads (otma; Proteintech) and incubated for 1 h while rotating at 4°C. Beads were washed three times in wash buffer (10mM Tris-HCl p=7.5, 150mM NaCl, 0.05% NP-40, 0.5 mM EDTA) and boiled in 2x SDS-loading buffer. Elution of HaloTag and HA-InlC was analyzed by western blotting using mouse anti-HA, mouse anti-Halo (28A8; Proteintech), and mouse anti-GAPDH.

### Statistics

The statistical analysis and significance are noted in the figure legends for each respective graph. Data was determined to be significant when P<0.05 for the indicated statistical test. All statistical analyses were performed in GraphPad PRISM 9.

## Contributions

Conceptualization: P.J.W., R.L.L. Methodology: P.J.W., R.L.L. Investigation: P.J.W., M.W.S., R.L.L. Writing the original manuscript: P.J.W., R.L.L. Revisions and editing manuscript: P.J.W., M.W.S., R.L.L.

## ACKNOWLEDGEMENTS

We thank Fabian Schutle and Brooke Linnehan at the Whitehead Institute Proteomics Core Facility and Richard Schiavoni at the Koch Institute Biopolymers & Proteomics Core Facility for experimental support. We would also like to thank Dr. Darren Higgins (Harvard Medical School, Cambridge, MA) for the *Listeria ivanovii* strain used in this study. We would also like to thank Brandon Sit and Jane Lodwick for critical feedback on the manuscript. P.J.W is a Damon Runyon Fellow supported by the Damon Runyon Cancer Research Foundation (DRG 2459-22). We would also like to acknowledge support from the National Institute of Health to R.L.L (R01GM141025 and R01AI155489).

## SUPPLEMENTARY FIGURES

**Figure S1.**
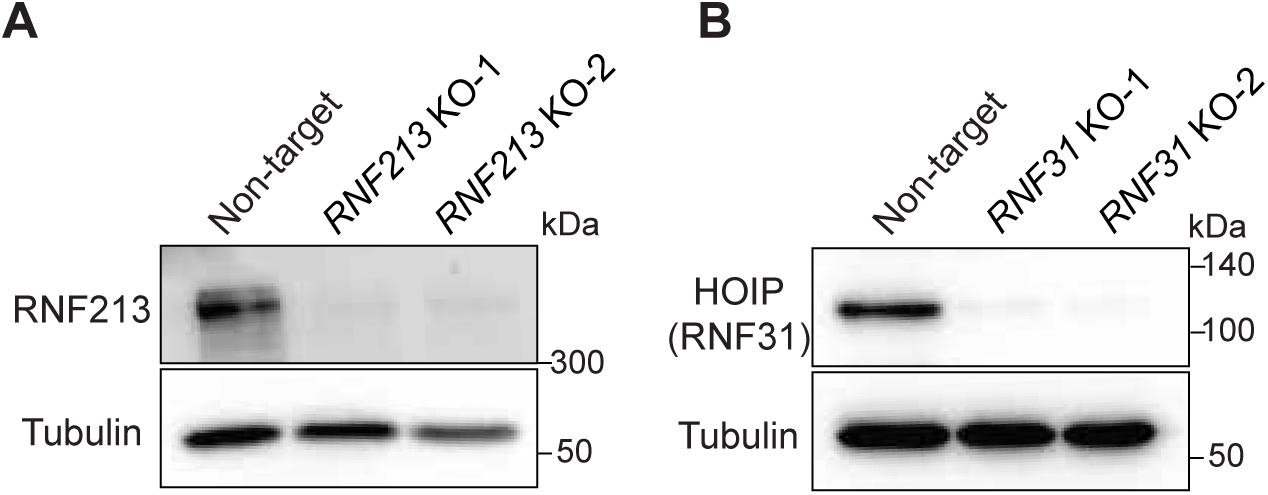
Western blot analysis of *RNF213* and *RNF31* knockout cells. **(A)** Western blot analysis of RNF213 from *RNF213* KO and non-target A549 cell lines. (**B**) Western blot analysis of HOIP (*RNF31*) from *RNF31* KO and non-target A549 cell lines.

**Figure S2.**
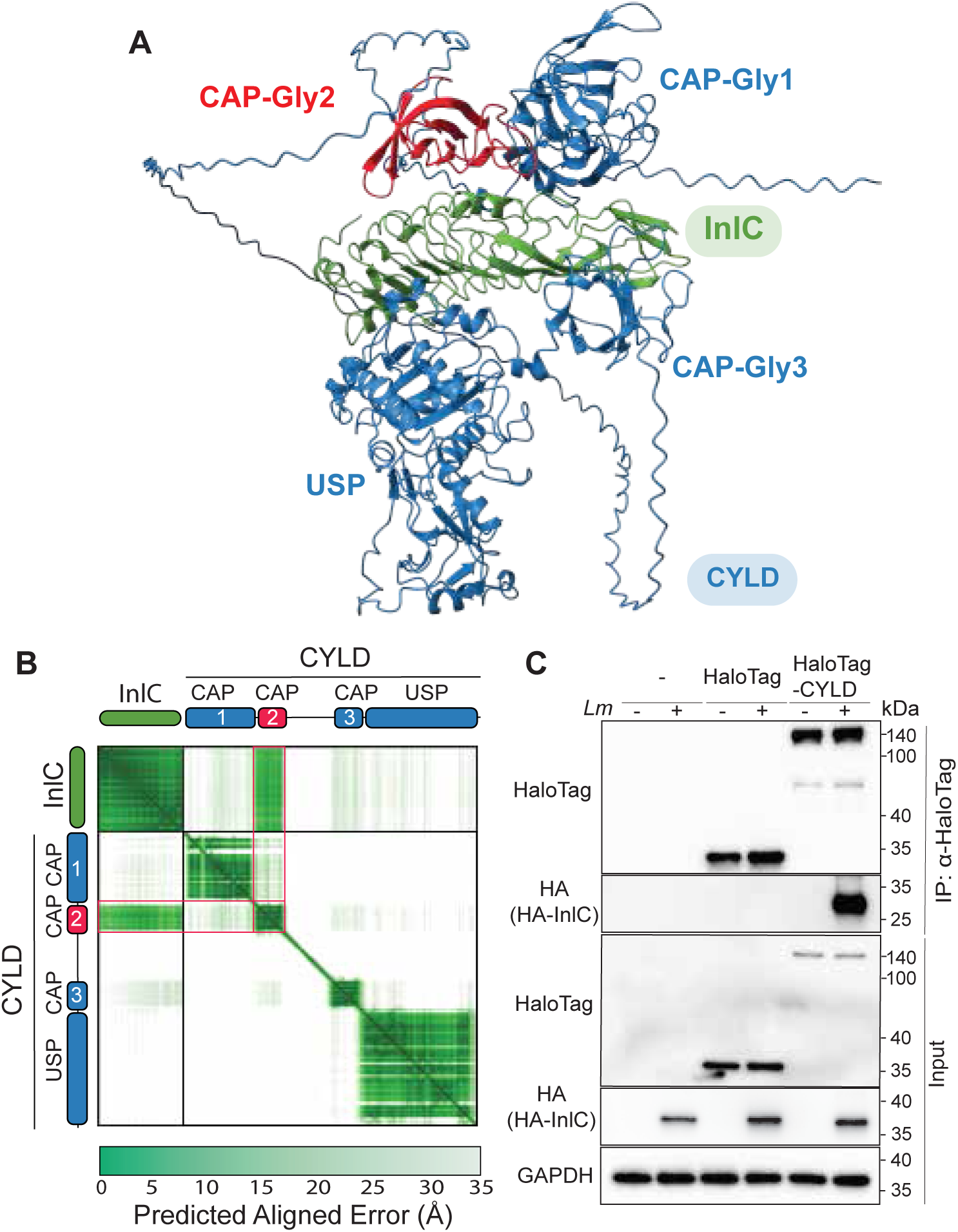
Alphafold structural prediction of InlC-CYLD binding. **(A)** Alphafold structural prediction model of InlC (green) and CYLD (cyan) predicts InlC binds to the second CAP glycine domain (CAP-Gly2, red). Additional CAP-gly domains and ubiquitin specific protease (USP) domains are indicated in blue. (**B**) ESMFold Error Plot displaying the Predicted Aligned Error (PAE) for the InlC-CYLD predicted structure from (A) with ipTM = 0.75. Low PAE (dark green) indicates high confidence in the relative position of the indicated residues. The top right square shows the PAE plot for InlC, and the bottom square displays the PAE plot for CYLD. CYLD plot identifies low PAE for the known CAP-Gly (CAP) and the USP. Side panels highlight PAE for the interactions between CYLD and InlC. High confidence interaction (low PAE) is suggested between InlC and the second CAP-Gly of CYLD. (**C**) Western blot analysis following Halo-Trap pulldowns from Δ*inlC* + InlC-HA *L. monocytogenes* infected A549, A549-HaloTag, or A549-HaloTag-CYLD cell lines. Data are representative of two independent experiments.

**Figure S3.**
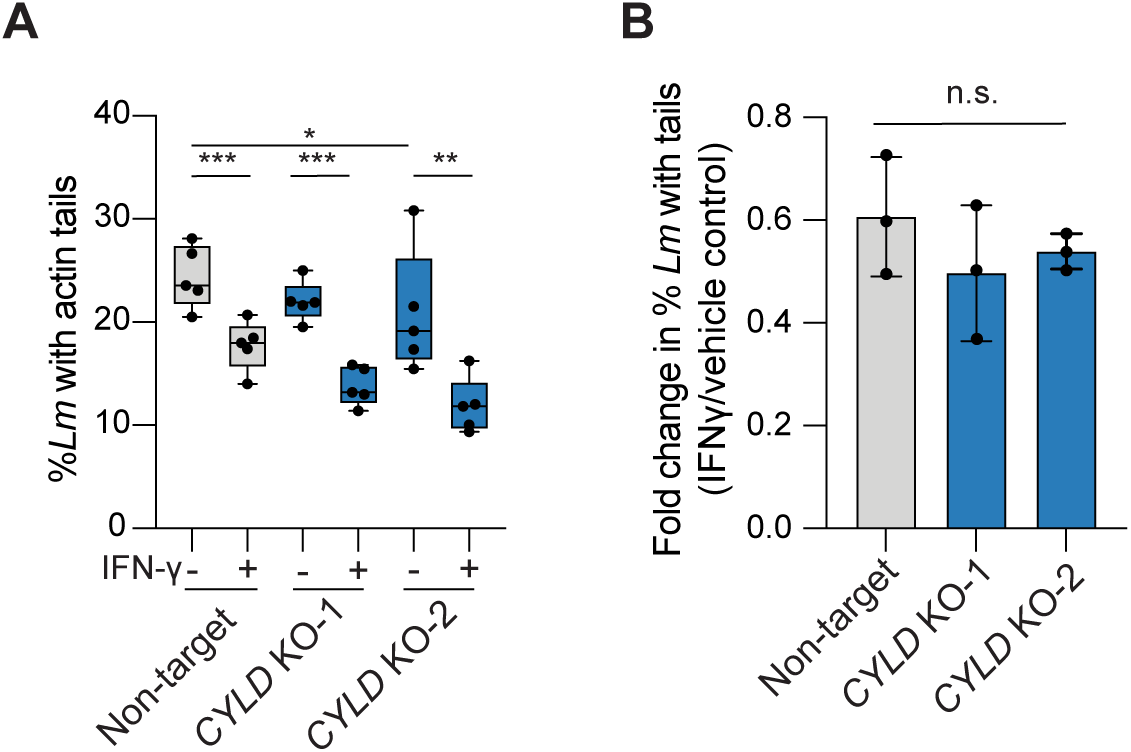
CYLD restriction of actin tail formation is not enhanced in response to IFN-γ. **(A)** Percentage of total *Lm*Dasher with actin tails in *CYLD* KO or non-target cells at 5 hpi, with and without IFN-γ. Data are representative of the percentage of indicated bacteria per FOV displayed as mean ± SD from three independent experiments; 170 > bacteria were counted per FOV in each experiment. (**B**) Fold change of the percentage of wild-type *Lm*Dasher with actin tails between IFN-γ treatment and vehicle control from (**A**). ANOVA using Tukey’s multiple comparison post-hoc test was performed on the averages from each individual experiment to determine significance \**P*<0.05, \*\**P*<0.001, \*\*\**P*<0.0001.

**Figure S4.**
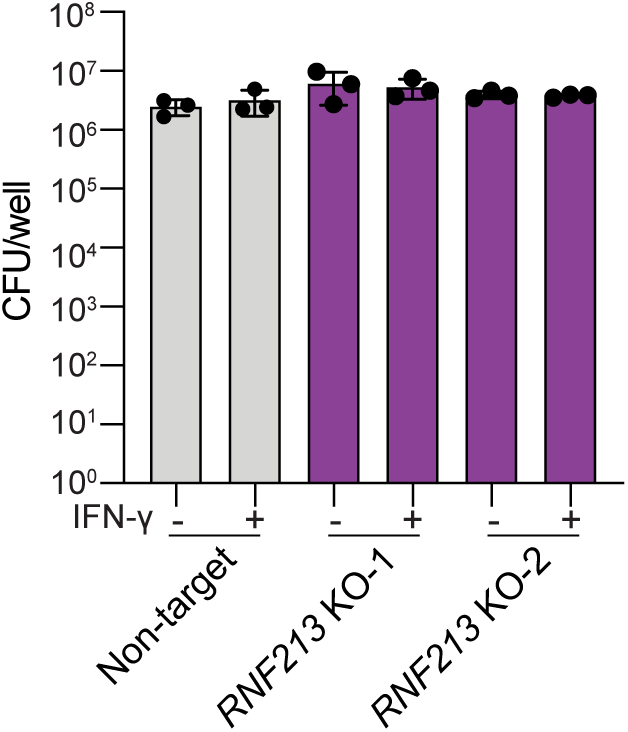
Loss of RNF213 does not impact bacterial burdens. Enumerated colony-forming units (CFU) per well of *L. monocytogenes* from infected *RNF213* KO or non-target A549 cell lines, with and without IFN-γ. Data representative of three independent experiments.

**Figure S5.**
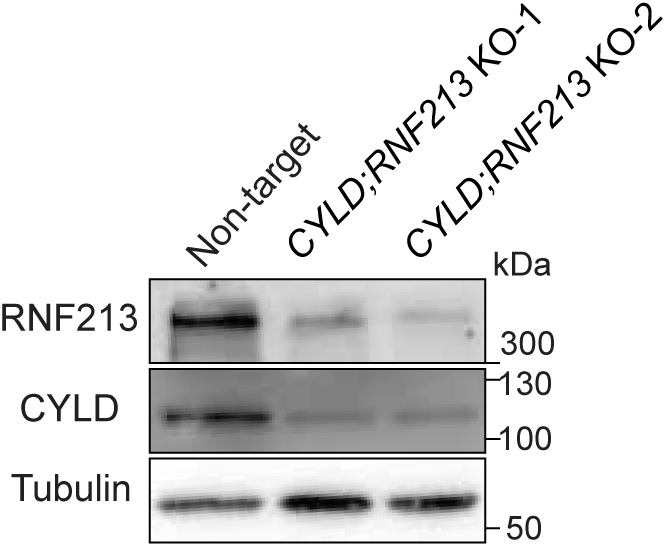
Western blot analysis of RNF213 and CYLD double knockout cells. Western blot analysis of RNF213 and CYLD from *RNF213;CYLD* double KO and non-target A549 cell lines.

## SUPPLEMENTARY TABLES

**Table S1. Surface display constructs designed from known *L. monocytogenes* surface proteins.**

**Table S2. Proteins enriched from streptavidin pulldowns of wild-type and Δ*actA Lm*NTurboID infected A549-CTurboID-V5-NES cells.**

**Table S3. Strains and plasmids used in the study.**

**Table S4. gRNA sequences for EnAsCas12a knock-out cell lines**

